# Accurate predictions of individual differences in task-evoked brain activity from resting-state fMRI using a sparse ensemble learner

**DOI:** 10.1101/2021.08.19.456783

**Authors:** Ying-Qiu Zheng, Seyedeh-Rezvan Farahibozorg, Weikang Gong, Hossein Rafipoor, Saad Jbabdi, Stephen Smith

**Affiliations:** Wellcome Centre for Integrative Neuroimaging, FMRIB, Nuffield Department of Clinical Neurosciences, Oxford, UK

**Author notes:** These authors contributed equally to this work (Saad Jbabdi, Stephen Smith).

## Abstract

Modelling and predicting individual differences in task-evoked FMRI activity can have a wide range of applications from basic to clinical neuroscience. It has been shown that models based on resting-state activity can have high predictive accuracy. Here we propose several improvements to such models. Using a sparse ensemble leaner, we show that (i) features extracted using Stochastic Probabilistic Functional Modes (sPROFUMO) outperform the previously proposed dual-regression approach, (ii) that the shape and overall intensity of individualised task activations can be modelled separately and explicitly, (iii) training the model on predicting residual differences in brain activity further boosts individualised predictions. These results hold for both surface-based analyses of the Human Connectome Project data as well as volumetric analyses of UK-biobank data. Overall, our model achieves state of the art prediction accuracy on par with the test-retest reliability of tfMRI scans, suggesting that it has potential to supplement traditional task localisers.

## INTRODUCTION

Studying individual differences in brain activity and how they relate to cognitive and genetic traits is an important area of research in basic and clinical neuroscience. Traditionally, functional Magnetic Resonance Imaging (fMRI) analysis has primarily been concerned with group-average inference. While averaging data across individuals substantially improves signal-to-noise (SNR) ratio and has proved fruitful in identifying common patterns across subjects, this approach treats unexplained individual variations as noise, discarding unique attributes of brain activity specific to a particular subject. Individual variations in neuroanatomical or functional activity often carries valuable information. For example, if a small number of subjects in a large cohort has a rare disease, an indiscriminate data reduction prior to the analysis will very likely obfuscate this information.

The rapid development of cutting-edge neuroimaging techniques in recent decades has led to substantial improvements in the reliability and validity of blood-oxygen-level-dependent (BOLD) measurements, providing an unprecedented opportunity to investigate individualised patterns of brain activity. Moreover, emerging “big data” projects, such as the Human Connectome Project (Van Essen et al. 2013) and UK Biobank (Sudlow et al. 2015), have collected multi-modal neuroimaging data on very large samples, enabling researchers to more closely examine individual variations in neuroanatomical patterns and functional activities with enhanced statistical power. Among previous fMRI studies of individual variabilities, an active line of research focuses on understanding how individual brains vary in response to external cognitive tasks. Following the work of (Tavor et al. 2016; Cole et al. 2016), a number of studies (Jones et al. 2017; Cohen et al. 2020; Ellis and Aizenberg 2020; Dohmatob et al. 2021) have since shown that spatial patterns of task-evoked activation form a stable trait marker, encoded in resting-state brain activity, i.e., in the absence of any explicit task. In contrast to previous studies that mostly rely on correlation analysis (in-sample inference) to investigate individual differences, these works adopt predictive frameworks that allow for out-of-sample inference and greatly improved generalisability of these investigations of individual variability.

Why is task-free fMRI predictive of task-evoked activation? Previous studies have suggested that resting-state networks and task networks may share the same intrinsic architecture (Smith et al. 2009; Cole et al. 2014; Krienen et al. 2014; Cole et al. 2016; Elliott et al. 2019). Therefore, a reasonable corollary is that resting-state heterogeneity should inform on variability of task-evoked brain activity. Typically, resting-state data are summarised as spatially continuous parcels distributed across the brain (Beckmann and Smith 2004; Calhoun et al. 2008; Van Den Heuvel et al. 2008; Calhoun et al. 2008; Bellec et al. 2010; Yeo et al. 2011; Craddock et al. 2012). These spatial maps are often referred to as “functional modes”, characterising functionally unified sub-processes underlying human cognition. Among the approaches of finding functional modes to predict task-fMRI, dual-regression (Beckmann et al. 2009; Filippini et al. 2009) is a widely-used algorithm, showing ability to predict individual idiosyncrasies in their response profiles (Tavor et al. 2016; Cohen et al. 2020; Dohmatob et al. 2021; Ngo et al. 2021). Although these previous attempts have successfully characterised individual-unique patterns of task-evoked brain activity, there are a few limitations yet to be accounted for. For example, these approaches focused on cortical regions and relied on pre-determined brain parcellations to extract predictors. Compared with models that take in global features without the need to *a priori* parcellate the brain, this introduces more free-parameters thus may increase the risk of over-fitting. Furthermore, these approaches did not attempt to explicitly model cross-subject variability of the rest and task states *per se*, and thus may be sub-optimal to capture cross-subject variations. In contrast, (Ngo et al. 2021) introduced a contrastive loss in combination with the common loss to maximise inter-individual differences. However, in practice, such loss functions are often non-convex and may have complicated behaviours (e.g. multiple local minimum) rendering optimisation difficult. To fully account for the inter-individual variations, an alternative is to explicitly train on residualised data, i.e., residuals where group-average information has been regressed out. The data obtained in this way has minimal shared variance with the group-level information, thus serves as a cleaner description of individual-level differences.

Here we propose a framework that explicitly models individual variations in task-evoked brain activity using the resting-state variability, the latter profiled by a set of common spatial modes derived from a recently developed technique, Stochastic Probabilistic Functional Modes (sPRO-FUMO). We show that, consistent with previous studies (Harrison et al. 2015; Harrison et al. 2020; Farahibozorg et al. 2021), sPROFUMO provides better sets of “bases” (later referred to as PFMs) to reconstruct the variations in task-evoked activation patterns than the widely used dual-regression. Additionally, we show that an ensemble learner that combines global and local bases has improved capacity of not only reproducing typical activation patterns but also preserved patterns unique to individuals. We demonstrate that modelling of individual-level task contrast maps comprises the modelling of two separate sources of variability, shape of activations and the overall activation strength. Considering these two aspects separately in task prediction is at least as effective as or even more desirable than simply modelling the original task contrast maps. Furthermore, the proposed model can recapitulate the spatial patterns of inter-individual variability, recovering regions that are more variable at the group-level. The model achieves state of the art prediction accuracy for both datasets, and is also on par with task test-retest reliability. These results demonstrate the potential of resting-state features to reproduce task-fMRI features, and serve as a supplement to task localisers in pre-surgical plannings.

## MATERIALS AND METHODS

### UK Biobank data

UK Biobank (UKB) is a large national project that collects a wide range of health-related measures for over 500,000 subjects, initially aged between 40 and 69. We used the resting-state and task functional MRI data from a total of 17,560 subjects. The acquisition parameters and processing details can be found in (Miller et al. 2016; Alfaro-Almagro et al. 2018). Briefly, all resting-state fMRI scans were acquired with identical scanners (3T Siemens Skyra) with a TR of 735ms for a total of 490 time points for each individual. After the initial preprocessing, the data were ICA-FIX cleaned to remove structured artefacts (Salimi-Khorshidi et al. 2014), and then registered to the standard MNI space. Next, each individual’s resting-state 4D time series were further spatially smoothed with a Gaussian kernel of sigma 3mm. The task used is the Hariri faces/shapes “emotion” task (Hariri et al. 2002; Barch et al. 2013), scanned and processed under the same protocols as the resting-state data (except that the task-fMRI data is not ICA-denoised). Individual as well as group-average activation z-statistic maps of three contrasts (faces, shapes, and faces-shapes) were estimated from the task fMRI scans using FEAT (Woolrich et al. 2001; Woolrich et al. 2004). Additionally, 473 subjects in this 17,560 subset received second-time scanning (mean test-retest-interval 2.25 years, std 0.12). These second-time scans provided test-retest reliability scores as a benchmark for our model performance.

### Human Connectome Project data

We used the MSMAll-registered data provided by the Human Connectome Project (HCP), S1200 Release (https://www.humanconnectome.org/study/hcp-young-adult). Details on the acquisition protocols and processing pipelines can be found in (Van Essen et al. 2013; Glasser et al. 2013; Robinson et al. 2014). Resting-state and task fMRI data from 991 subjects, aged 22 to 35 years, were used in the analysis. Each individual had four runs of resting-state scans with a TR of 0.72s for a total of 1,200 time points per run. The data were ICA-FIX denoised to remove the effect of structured artefacts automatically, then resampled onto the “32k_fs_LR” grayordinates space and minimally-smoothed by 2mm FWHM. All subjects were MSMAll-registered to improve functional and structural alignment (Robinson et al. 2014). To further increase the signal-to-noise ratio, an additional smoothing of 4mm FWHM was applied to the MSMAll-registered data (with subcortical structures smoothed within parcel boundaries, and cortical data smoothed in 2D on the surface) using the Connectome Workbench (https://www.humanconnectome.org/software/connectome-workbench). The task fMRI scans were acquired and pre-processed in the same way (though without FIX). We used the MSMAll-registered individual and group-average contrast maps with 4mm FWHM smoothing in the analysis, including 47 contrasts across seven task domains (Barch et al. 2013).

Similarly to the UKB dataset, we used the HCP retest scans as the reliability benchmark for the predictions. Among the 991 subjects, 43 have received second-time scanning under the same 3T imaging and behaviour protocols with test-retest-interval ranging from 18 to 328 days (mean 134.78; std 62.49).

### Generation of resting-state functional modes

We used resting-state functional modes to predict individual task-fMRI. Functional modes are typically modelled as parcel-like spatial configurations of unified functional processes distributed across brain, each characterised by a summary time course that captures mode activity over time. Here we explored two approaches of generating individual resting-state modes, group-ICA followed by Dual-Regression (DR-ICA) and Stochastic Probablistic Functional Modes (sPROFUMO). DRICA is a conventional group-average algorithm to estimate individual “versions” of group-level spatial configurations, using a set of common spatial modes as templates (Nickerson et al. 2017). In DR-ICA, group-PCA was carried out on each dataset (UKB and HCP) by MELODIC’s Incremental Group-PCA (Smith et al. 2014) on the resting-state time series of all subjects (temporally demeaned and variance normalised), producing 1,200 weighted spatial eigenmaps for UKB and 4,500 eigenmaps for HCP. These eigenmaps were subsequently fed into ICA using FSL’s MELODIC tool to generate group-ICA spatial maps at multiple ICA-dimensions (i.e., the number of distinct ICA components). To obtain dual-regression maps for a specific subject at a given ICA-dimension *k*, we first regressed the corresponding *k*-dimensional group-ICA spatial maps into the individual 4D time-series data, yielding a set of *k* time courses per subject. The resulting time courses were subsequently regressed into the same 4D time-series, generating *k* dual-regression spatial maps for each subject. However, a major limitations of DR-ICA is that it only allows unidirectional flow between group and individuals, i.e., the estimated individual modes cannot in turn drive the refinement of group-average modes, and may have limited ability to cope with individual deviations from the group-average (Bĳsterbosch et al. 2018; Bĳsterbosch et al. 2019). A recently developed technique, sPROFUMO, uses a Bayesian model that simutaneously estimates functional modes both at group- and individual-level, and is scalable to large datasets (Farahibozorg et al. 2021). In sPROFUMO, individual resting-state time-series are factorised into a set of spatial modes and their summary time courses (one per mode), together with the time course amplitudes. The group-level parameters constrain the estimation of (the posteriors over) individual-level parameters, of which the posterior evidence is accumulated across individuals to in turn infer the group-level parameters. The bidirectional information flow between the group and individuals aims to result in improved subject-specific spatial alignments. Below, we refer to the resting-state feature maps as either DR-ICA maps or PFMs depending on the approach used to derive them.

### Residualisation of the resting-state and task contrast maps

Our aim is to derive a model that can predict task activation in individuals given their resting state modes. One of the innovations in this paper is to try to explicitly capture individual variations in our model. We propose that training and evaluating the model on residualised data (i.e., data and features where the group-averaged maps have been regressed out) would be of more value than training a model on the original resting-state and task contrast spatial maps.

To understand this, consider each individual task contrast map. It can be decomposed into the sum of the group-averaged map (scaled by some factor) and a spatial residual map specific to the individual. The resting-state feature maps can also be similarly decomposed. Once a model has been trained, the correlation between the model prediction and the task map of a test subject is composed of four terms: (i) the correlation between the group averaged task map and the subject’s scaled version of that map (this correlation is close to or equal to one, scaled by amplitude), (ii) the correlation between the residual maps for prediction and test data, and two cross terms (iii) the correlation between group-average and residual target, (iv) the correlation between group-average and residual prediction. The cross term (iii) will disappear because of the orthogonality between group-average and residuals, and the cross term (iv) is very close to zero in practice. Hence, only (i) and (ii) significantly contribute to the overall correlation, while (i) can bias the prediction towards the average subject. By residualising both the target task maps and the resting-state feature maps with respect to their group averages, we can remove this bias and better model individual variations.

Hence, we built the model entirely on residualised data. The residualised resting-state functional modes and the residualised task contrast maps are also referred to as “resting-state variation maps” and “task variation maps” in the remainder of the paper. To residualise the resting-state data, each of the *k* group-average (across training subjects) ICA spatial maps was regressed out from the corresponding individual DR-ICA maps for all subjects (i.e., a one-variable linear spatial regression per subject per dual-regression map), and similarly, each sPROFUMO group-level spatial map was regressed out from the same mode’s individual-level sPROFUMO spatial maps (PFMs). These residualised spatial maps represent individual variations in resting-state activity, serving as features to predict individual variations in task-fMRI.

The task activation maps were residualised similarly for each individual, to give task variation maps. For a given task contrast, the group-average activation map was regressed out from the individual contrast maps (i.e., a simple linear regression per subject per contrast, with the group-average activation map as the regressor). These task variation maps describe individual differences in task-evoked brain activity that deviate from the typical activation patterns. Therefore the mapping between the rest and task states is entirely based on the variations rather than on the original resting-state and task-evoked activity. See Figure 1 b for an illustration of residualisation.

**Figure 1.**
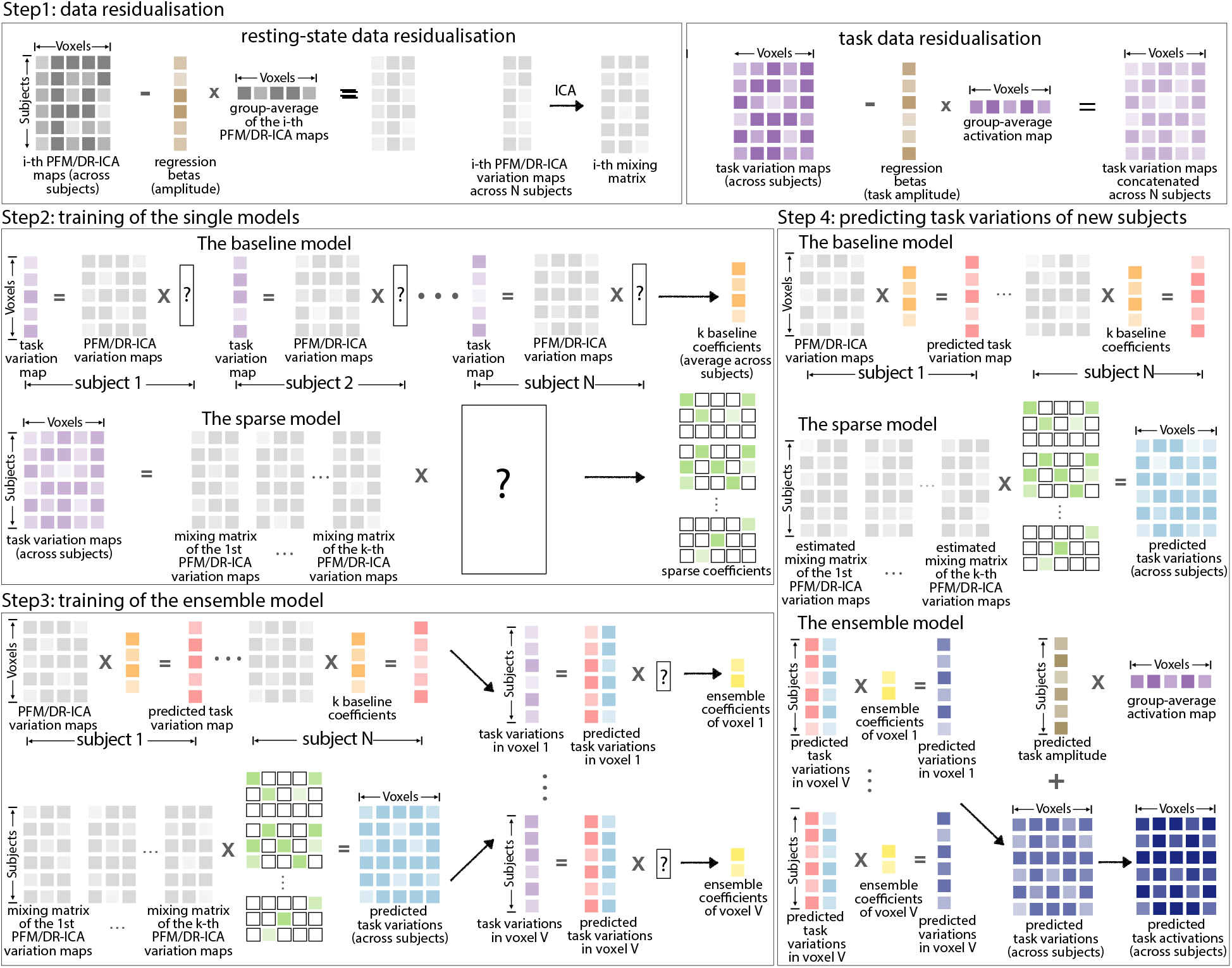
An illustration of the model. **Step 1.** Residualisation of the resting-state models and task contrast maps. The residualised resting-state maps were further ICA-reduced as the input of the sparse model. **Step 2.** Training of the baseline and sparse model: per training subject, the baseline model yielded *k* reconstruction coefficients (one coefficient per map), which were averaged across subjects as the final baseline coefficients (orange). Next, the resting-state variation maps and the task variation maps were concatenated across subjects accordingly and then reduced to lower dimensions via ICA. The sparse model was trained on the (ICA-reduced) across-subject variation matrices to give the sparse regression coefficients (green). **Step 3.** the estimated baseline coefficients and sparse coefficients were applied to the training subjects to get the baseline-model-fitted (pink) and sparse-model-fitted (blue) task variation maps. Next, for each voxel across subjects, we estimated the ensemble coefficients (yellow) by fitting another linear regression model with the baseline-model-fitted activations and the Lasso-fitted activations (in the corresponding voxel) as the two regressors. **Step 4.** These three sets of coefficients were finally applied to the test subjects to make new predictions (navy).

Finally, to compare this residualised model with the model trained on the original resting-state and task contrast maps (i.e., un-residualised), the task activations are predicted as a combination of the modelled variations and the average activation patterns. For both the rest and task data, we record the regression parameters as part of regressing out group-mean maps; these measures of overall “amplitude” are used later in the work, described below, and can of course be used (multiplied by the group-mean maps) to add the group-mean contribution back in where desired.

### The ensemble learner

Our overall ensemble approach combines two separate models, “baseline” and “sparse”. We start by describing these two individual models, and then go on to describe the ensemble method.

#### The baseline model

The baseline model assumes that, for a given task contrast, the individual task variations (i.e., residualised activation maps) can be represented by a linear combination of the variations in resting-state functional modes (i.e., residualised DR-ICA maps or PFMs). In this sense, the resting-state modes serve as a set of “bases” that span the task space. To obtain the reconstruction coefficients for each subject, we regressed the subject’s resting-state bases into its spatial activation map (i.e., a multiple-regression per subject per task contrast, with resting-state variation maps as regressors and the task variation map as response). More specifically, suppose the number of voxels is *V*, and each individual has *k* number of bases (i.e., there are *k* group-average ICA spatial maps); to find the reconstruction coefficients of a specific task contrast map **y** *_j_* (a *V* × 1 vector) for a given subject *j*, we regress the given subject’s *k* resting-state variation maps, denoted by a *V* × *k* matrix 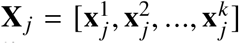 for *i* = 1, 2, …*k*, into the task variation map **y** *_j_* of this subject. As a standard linear regression problem, the reconstruction coefficients ***β*** *_j_* (a *k* × 1 vector) of subject *j* minimise the following loss function

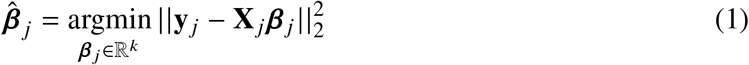

for *j* ∈ *S*, where *S* is the set of training subjects. The estimated reconstruction coefficients 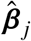 are given as 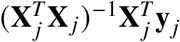, where **X** *_j_* is subject *j*’s resting-state variation maps. These coefficients were averaged across the training subjects to give the final estimates of the reconstruction coefficients, i.e., 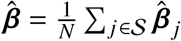, where |*S*| = *N* is the number of training subjects.

To predict the activation map of an unseen subject *l* in a test set 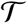, we applied the reconstruction coefficients averaged from the training set to the subject’s own resting-state variation maps **X***_l_*, i.e.,

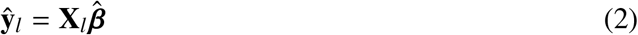

for 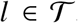. Note that the baseline model is different from (Tavor et al. 2016; Cohen et al. 2020; Dohmatob et al. 2021) in two ways. First, their models were primarily local, i.e., one linear regression per brain region, rather than a global linear regression for the whole brain. Second, with the group-average content regressed out from both the resting-state dual-regression maps and task activations, our baseline model aims to establish linear relationships between the variations of the two states (relative to the group-average) rather than the original resting-state and task activity (which is possibly dominated by the group average).

#### The sparse model

The baseline model has a few limitations. First, it has very few free parameters, resulting in one reconstruction coefficient per basis, which is then pooled (averaged) across all subjects. Crucially, each feature (spatial map) is associated with a single regression coefficient, regardless of which part of the brain is being modelled. Second, the coefficients learned from each training subject are estimated separately, which ignores patterns of between-subject variations. We re-formulated the problem in a more flexible, highly-parameterised framework, referred to below as the sparse model, with appropriate regularisation techniques to protect against too much flexibility.

First, to create feature maps that contain information of cross-subject variability, each of the *k* resting-state variation maps is first concatenated across the set of training subjects *S*, yielding one *N* × *V* resting-state matrix per group-average spatial map, with a total of *k* such matrices. Denoting the *i*-th matrix 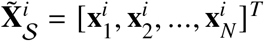, where 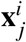 is the *i*-th resting-state variation map of subject *j*, we then dimensionality-reduce these matrices into a set of *d* components using ICA: 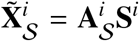 for *i* = 1, 2, …, *k*. Following this decomposition, 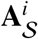 is an *N* × *d* mixing matrix and **S**^*i*^ is a set of *d* independent components representing common spatial variations across the training subjects *S*. The mixing matrices of each ICA contains “coordinates” of each individual in the resting-state space spanned by these common modes, providing profiles of the resting-state variabilities of these individuals. The *k* mixing matrices are concatenated to give a single reduced variation matrix 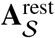 as the final predictors, where 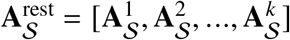 is an *N* × *dk* matrix.

Likewise, the task variation maps (residualised activation maps) are concatenated across the training subjects *S*, resulting in an *N* × *V* task variation matrix **Y**_*S*_ = [**y**_1_, **y**_2_, …**y***_N_*]^*T*^ per contrast. The reduced resting-state variation matrix 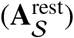 will be used to predict the concatenated task variation matrix. Under this formulation, the model has a large number of potential predictors. To prevent over-fitting, we enforce sparsity on the prediction regression coefficients, to enable selection of the subset of features that are most desirable for prediction. In addition, given that predictions made on the original task matrix are not only computationally expensive but also involve many redundant and noisy features (which will likely compete with the “real” features in the training), we also consider to similarly decompose the task matrix into a set of *p* independent components, i.e., 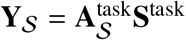, where 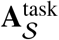 is the *N* × *p* mixing matrix, and **S**^task^ is the set of *p* independent components. Thus, both the features matrix 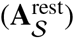 and the regression target used in training (**Y**_*S*_) are sparse, low-rank versions of their original versions (through ICA), and contain information on individual variations (through concatenation of subjects).

To find the sparse coefficients **W**, we solve the following regularised regression problem on the ICA-reduced task matrix 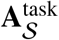

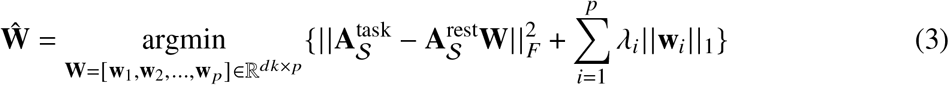

or on the original task maps **Y**_*S*_

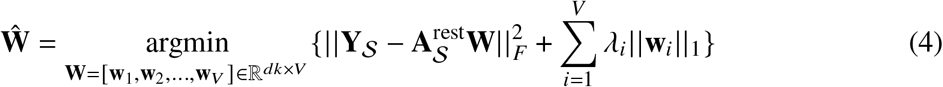

where the Lasso penalty is univariately applied to columns of **W** (with different hyper-parameters to allow differential amounts of regularisation), encouraging it to be element-wise sparse. Note that an alternative way of introducing sparsity is to use an *L*_1,2_ penalty on **W** that enforces row-wise sparsity, as commonly applied in the grouped Lasso and the multivariate Lasso. That strategy would permit simultaneous use of all outputs to estimate a sole regularisation parameter. It implicitly assumes that predictions of different outputs (columns of 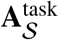 or **Y**_*S*_) tend to require the same set of features. This underlying assumption of row-sparsity penalty is not very appropriate and tends to require heterogeneous feature selection. Other alternatives that simultaneously use all outputs include Partial Least Squares (PLS), Canonical Correlation Analysis (CCA), and their variants, as well as a range of multi-task learning approaches. Given that multi-task learning approaches with sparsity regularisations usually have more complex behaviours than the pure Lasso, we simply choose the Lasso penalty, which is also particularly easy to parallelise across columns of 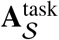 or **Y**_*S*_ (i.e., across task voxels).

To predict task variation maps for a set of unseen subjects, denoted by 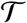, we first need to translate the subjects’ resting-state variations into the subspace spanned by the resting-state common modes (decomposed from the training subjects). This is conducted by regressing each across-subject basis matrix onto the corresponding set of resting-state common modes. Again, suppose the *i*-th (across-subject) resting-state variation matrix of the test subjects is denoted by an *n* × *V* matrix 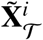, where 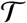 is the test set, and 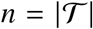 the number of test subjects. We seek to solve the linear regression problem

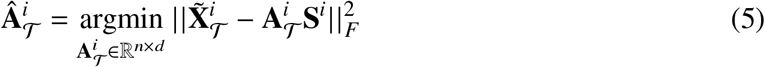

where 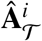 is the estimated “mixing matrix” of the *i*-th resting-state variation matrix across the test subjects 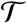, and **S**^*i*^ is the independent components calculated from the training subjects for *i* = 1, 2, …, *k*. Next, the sparse coefficients 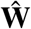, estimated via (3) or (4), are applied onto the concatenated variability profiles 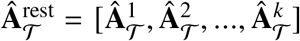 (an *n* × *dk* matrix), to give predictions for the set of unseen subjects 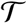

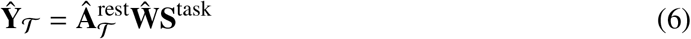

if 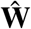 is solved via (3) or

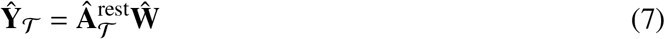

if 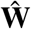 is solved via (4).

This completes the specification of the sparse model. To summarise the approach, we use concatenation of training subjects to incorporate information on subject variability in the training, we apply ICA to sparcify this data to help with fitting, and we employ further regularisation via the Lasso cost function on the regression coefficients. For UKB, we chose to reduce each across-subject resting-state matrix to 3,000 independent components and the task matrix to 4,000 independent components (however, reducing the resting-state and task matrices to 1,000 independent components yields comparable results, see Figure S5 a). For HCP, in contrast, we chose to reduce each resting-state matrix to its full rank (i.e., number of the training subjects, around 900 in each fold) but kept the original spatial dimension of each task matrix (i.e., no ICA on the task matrix), which yielded the best performance on a left-out subset (see Figure S5 b).

#### The ensemble model

A single model usually represents a single hypothesis space of the particular prediction problem. Although the single models may contain the hypothesis space already well-suited for a specific problem, combining multiple hypotheses allows for more flexible structures to exist between predictors and response variables, and can potentially improve model performance (again, as long as over-fitting is avoided through correct use of, for example, cross-validation or left-out data). Here the two single models are tailored to different aspects of the underlying hypothesis that variations in resting-state activity can inform task variations. As mentioned above, the baseline model treats the resting-state variation maps as a set of “bases” that spans the task variation map for each individual. It is obvious that the baseline model assumes that the mapping between resting-state and task space is within-subjects and thus ignores between-subject patterns of variations which might also be useful for the predictions. The underlying hypothesis of the sparse model captures a different aspect, though closely-connected with the baseline model hypothesis. With the variation maps reduced to the corresponding subspaces, the sparse model assumes that the “coordinates” of the subjects in resting-state space can be translated into their “coordinates” in task space.

Here we aggregate the predictions of each single model, to give the final prediction for unseen subjects using simple linear regression. Suppose 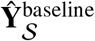 is the *N* × *V* baseline-model-fitted activations of the training subjects *S*, and 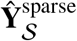 is the sparse-model-fitted maps. Particularly, we use 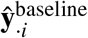 and 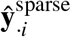 to denote the fitted activations in voxel *i* across subjects (i.e., each is an *N* × 1 vector). At the ensemble stage, we aim to find the coefficients for each constituent model by column-wisely fitting a simple linear regression on the task matrix of training subjects **Y**_*S*_, i.e.,

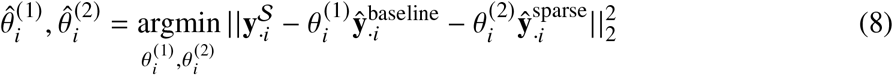

for the “true” activations in voxel *i* across the *N* training subjects, denoted by 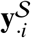, for *i* = 1, 2, …, *V*. The two coefficients, ^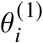^ and ^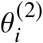^, will then be applied to the baseline-model-predicted and sparse-model-predicted maps to yield predictions of task variations for the unseen subjects. See Figure 1 for an illustration of model training and Table S2 for a summary of the notations.

For UKB, the ensemble model and its constituent base models were trained and tested on 17,560 subjects (3-fold cross-validation); for HCP, the models were trained and tested on 991 subjects (10-fold cross-validation). The hyper-parameter of the *L*_1_ penalty was optimised within each fold’s training data via nested cross-validation (3-fold). The other free parameters (e.g., the number of resting-state bases and the number of independent components in the sparse model) were determined on a different subset of 4,700 subjects for the UKB dataset (trained on 4,000 and tested on 700). Due to the limited number of HCP subjects, we randomly selected 10% of the HCP subjects and investigated how the choice of these parameters would affect the model.

#### The amplitude model

The amplitude model aims to predict the task activation amplitude for each individual (i.e., the beta coefficients from regression against the group-average activation map, as recorded during residualisation, one scalar value per subject per contrast) using the resting-state amplitude (i.e., the beta coefficients from regression of resting maps against the group-mean dual-regression maps, one scalar value per subject per basis). There are a few reasons for incorporating a separate amplitude estimation. First, one important source of individual variabilities in task-evoked activity is the (overall) activation amplitude. Explicitly predicting this information may help capture a different kind of individual variability that cannot be fully modelled by the aforementioned spatial models (indeed we would not expect to capture this from the residualised predictions). Second, the final predictions for test subjects are ideally given as a combination of modelled residual variations and the typical activation patterns. In order to recover the activation maps from the variation maps for each individual, the group-average activations are yet to be added back in appropriately, scaled by the activation amplitude of the specific individual. However, the activation maps of the test subjects are of course not seen during training. Therefore, we are not able to estimate the activation amplitude (i.e, betas from task residualisation) by simply regressing the group-average activations into the inidividual activations. As an alternative, the resting-state amplitude may be predictive of the overall activation amplitude (Figure S1 and S2) and thus may serve as a substitute for this information. The surrogate activation amplitude was generated as follows. Remembering that each subject has *k* resting-state amplitude values, corresponding to each of the *k* group-average spatial maps (i.e., one amplitude value per map): for a given contrast, a multiple linear regression model with the activation amplitude as the response and the resting-state amplitude as the predictors was trained across subjects (3-fold cross validation on UKB; 10-fold cross validation on HCP). These surrogate activation amplitude are subsequently applied to the predicted variation maps as the new beta coefficients, such that the re-scaled group-average effects can be added back in accordingly.

The other hypothesis about the overall activation amplitude is that it serves as another important source of individual variabilities. To explore this possibility, we also consider to incorporate the amplitude information into the ensemble stage to test whether it can further improve model performance. Given that the *k* resting-state amplitude values of the *k* sets of dual-regression maps are correlated (across subjects), we reduce the *k* amplitude features into a few principal components, the number of which are determined via cross-validation. These components are included in the ensemble model as additional predictors to predict each column (voxel) of **Y**.

### Measures of model performance

Assessment of model performance is primarily based on Pearson’s correlations between predicted maps and the actual maps (in subjects left out of the training process). Apart from the standard MNI152 brain mask applied at the beginning of all the analysis, we choose not to apply further thresholding of the resulting maps. Although further masking of the images may emphasise certain regions that are more of interest, the choice of thresholds can have a complex impact on evaluation and requires caution.

For a given task contrast, the predicted maps are correlated with the actual maps for all subjects, yielding a “subject by subject” correlation matrix, where the entry in the *i*-th row and *j*-th column corresponds to the correlation between subject *i*’s predicted map and subject *j*’s actual map. The mean of the diagonal elements measures the overall prediction accuracy, i.e., how well the model can reproduce the spatial patterns of activation for each subject, averaged across subjects. However, this measure cannot fully quantify model performance because the overall model accuracy can be boosted by simply reproducing the group-average activation, particularly when most subjects are “normal”, having activations patterns close to the group-average. Therefore, it is also important to make differentiated predictions, i.e., how well the model can capture atypical variations that deviate from the group-average activations. This necessitates measuring the extent to which, for a specific subject, the model can make predictions that are closer to the subject’s own activation maps than to the others. This is of course particularly relevant if doing non-residualised prediction.

The new evaluation measure is calculated as follows: after the correlation matrix (between predicted maps and the actual maps for all subjects) is normalised via Fisher’s transformation, for each subject, we calculate the difference between two values: (i) correlation between the subject’s predicted map and the subject’s actual map; (ii) mean of the correlations between the subject’s predicted maps and other subjects’ actual maps. The difference between (i) and (ii) provides a quantitative evaluation of the model’s capability of predicting individual differences distinct from the group mean. In the following text, the first measure is referred to as “prediction accuracy”, and the second one is referred to as “prediction discriminability”.

Additionally, we calculated the between-subject standard deviation map of the actual task variations (as a measure of inter-individual voxel-wise variability) and also of the predicted variations (as a measure of predicted variability) for each contrast. We then correlated the predicted variability maps against the actual variability map as a third measure of model performance. A higher correspondence between the two standard deviation maps indicates better ability to reproduce the spatial pattern of between-subject variability.

## RESULTS

### The ensemble model outperforms its constituent single models

To compare DR-ICA maps with PFMs, we chose the optimal dimensionality of each method, DR-ICA25 and PFM-50 for UKB, and DR-ICA50 and PFM-150 for HCP, respectively. The fact that PFM optimal dimensions were found to be higher than those of DR-ICA suggests that the former yielded more reliable functional modes particularly at higher dimensions (however, note that PFMs consistently outperformed DR-ICA across all dimensions. See Figure S3). In the baseline model, overall, most variation maps contributed to the predictions (Figure S4), suggesting that these resting-state variation modes did capture a significant proportion of the variance in task variation maps.

We found that the sPROFUMO modes had overall higher accuracy in predicting task variations than the DR-ICA maps, consistently across the baseline, sparse, and ensemble model (Figure 2). Compared with predictions based on DR-ICA, the biggest improvement introduced by sPROFUMO modes was evident from the baseline model, suggesting that sPROFUMO provides a fundamentally better set of resting-state bases to reconstruct task variations than DR-ICA. This corresponds with previous evidence that sPROFUMO better accounts for cross-subject misalignment and accommodates higher predictive power of population heterogeneity (Farahibozorg et al. 2021). Additionally, sPROFUMO modes also exhibited higher prediction accuracy for the sparse and ensemble model. Interestingly, the baseline and sparse model based on DR-ICA had very distinct performance on the two datasets. For HCP, the baseline model yielded higher prediction accuracy than the sparse model (Figure 2b, blue and green), while for UKB, this relationship was entirely reversed (Figure 2a, blue and green). However, introducing sPROFUMO modes as bases substantially enhanced prediction accuracy of the baseline model for UKB, making it tend to outperform the sparse model. Here we provide a possible explanation for this discrepancy. The two single models are tailored to different data scenarios. If the resting-state modes form the set of bases that do fundamentally have the ability to predict the task maps, then the baseline model should suffice, i.e., we don’t need the sparse model to emphasise specific spatial features. In practice, however, DR-ICA maps are not the perfect sets of individual “versions” of the group-average modes, containing many noisy voxels irrelevant to task-fMRI prediction. A major difference between the two datasets is that the UKB data we used to train the model is volumetric while the HCP data is grayordinates. As a consequence, there is more functional spatial variability (misalignment) in the UKB data (Coalson et al. 2018) and hence more “errors” in its individual dual-regression maps. In addition, HCP data is MSMAll-aligned and UKB is not. On the other hand, sPROFUMO better accounts for cross-subject misalignment and allows more fine-grained delineation of individual differences in resting-state data, thus it has improved ability to capture variations in task data. Furthermore, due to the shorter scanning sessions, the resting-state and task-fMRI scans in UKB have higher noise than in HCP, requiring additional benefits of identifying which voxels/spatial features are more desirable in the modelling. Hence, UKB requires greater spatial modelling complexity as well as greater spatial smoothing, provided by the sparse model (note that conducting ICA on the resting-state and task matrices across subjects in the sparse model may serve as a kind of de-noising).

**Figure 2.**
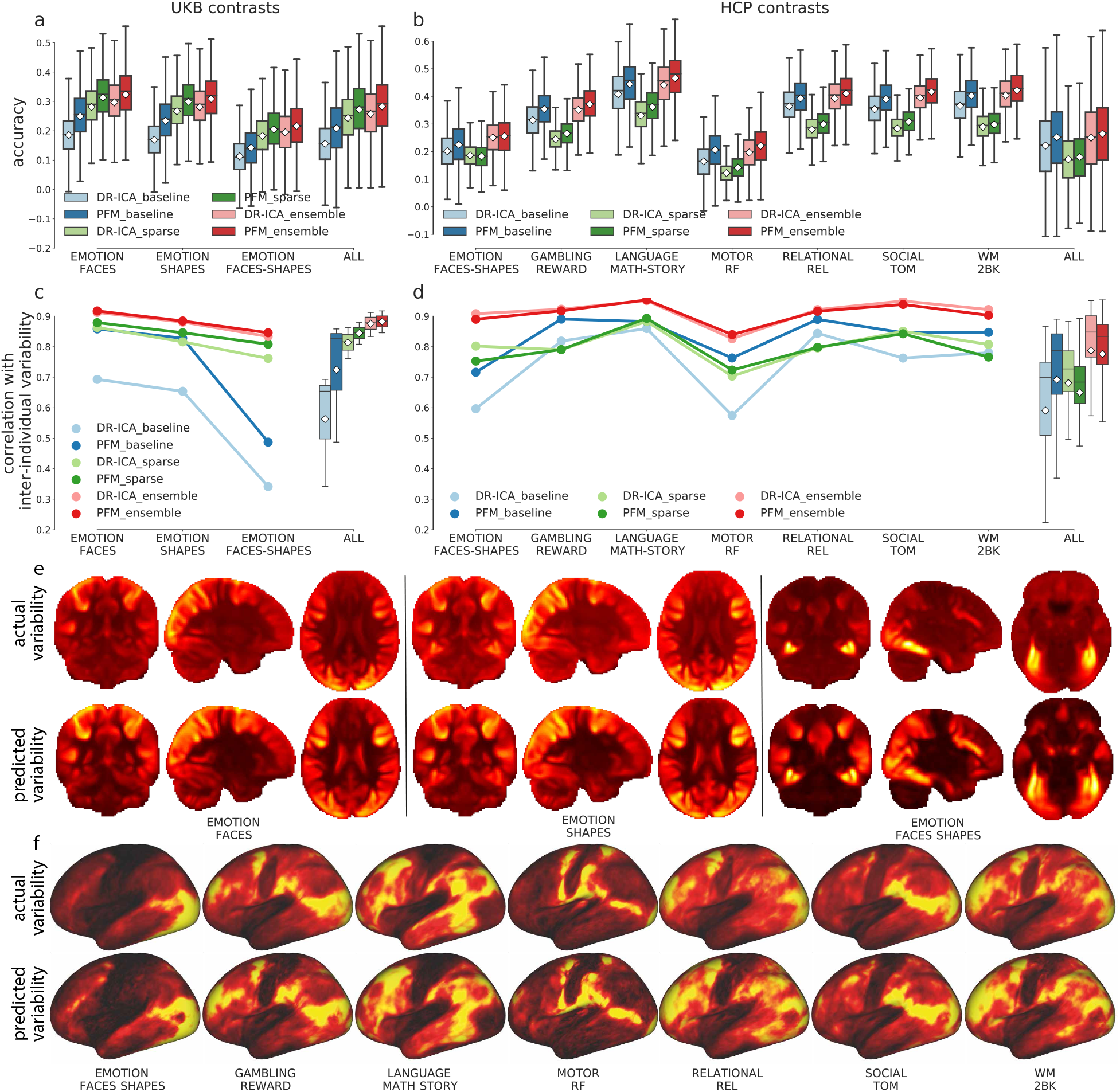
Prediction accuracy of the individual task variations and of the inter-individual variability. PFM better captures task variations than DR-ICA maps (dark colors vs pale colors); the ensemble model outperformed its constituent single models in predicting individual task variations and reproducing inter-individual variability patterns (blue, green, and red). (a) Prediction accuracy of the baseline, sparse, and ensemble models for 17,560 UKB subjects across the three contrasts, the last columns showing all contrasts pooled together. White diamonds show the means along with the boxplots showing the medians and quartiles. (b) Equivalent plots of 991 HCP subjects across seven representative contrasts, the last column showing all 47 contrasts pooled together. (c) Correlations between the predicted and the actual inter-individual variability maps calculated across 17,560 UKB subjects. Overall, ensemble trained on PFM yielded the highest correspondence with the inter-individual variability. (d) Equivalent plots across 991 HCP subjects. See Figure S7 for all HCP contrasts. (e) The actual (first row) and the predicted (second row) inter-individual variability across 17,560 UKB subjects of the three contrasts, shown volumetrically. Warmer colors indicate higher variability with the maximum normalised to 1. (f) The actual (first row) and the predicted (second row) inter-individual variability calculated across 991 HCP subjects of the seven representative contrasts, shown on the cortical surface.

For both datasets, overall, the ensemble model outperformed its constituent single models. Remember that the task variations are the residuals of regressing the group-average activations into the individuals, thus they are orthogonal to the group-mean by design. This also implies that these task variation maps have minimal overall cross-subject similarity, i.e., the spatial correlations between pairs of subjects fluctuate around zero. Therefore, the plots of prediction accuracy and of discriminability will look almost identical, because the predicted maps will have near-to-zero correlations with the maps of the other subjects, i.e., the off-diagonals of the (subject by subject) correlation matrices (between the predicted maps and the actual maps) are all close to zero (Figures S8 and S9).

In addition to predicting the individual variations in task activity, all three models could reproduce the spatial pattern of inter-individual variability (standard derivation maps across subjects) for both datasets (Figure 2c and d). Similar to the previous scenario, using sPROFUMO modes as bases improved the prediction of inter-individual variability for the baseline model on both datasets (Figure 2c and d, blue), corroborating the conclusion that sPROFUMO better aligns the subjects, refines the spatial details of cross-subject heterogeneity, and thus provides a better set of bases to reconstruct task variation space. In terms of the sparse and ensemble model, DR-ICA and sPROFUMO yielded comparable correspondence with the true inter-individual variability.

These actual and predicted (via the ensemble model) inter-individual variability maps are shown in Figure 2e and f. Regions of higher variability across subjects are those more involved in the corresponding task execution. For example, somato-sensory and motor regions are more variable across subjects in the motor contrasts; fronto-parietal regions exhibits higher variability in more cognitive contrasts; the visual areas tend to be more variable in general, for all contrasts. In summary, all three models are able to capture individual-unique activation patterns that deviate from the typical activation patterns as well as recapitulating the spatial pattern of inter-individual variability. In the subsequent analysis, we used PFM50 for UKB, and PFM150 for HCP. The subjects identification accuracy (i.e. the probability that predicted maps had the highest correlation with the subjects’ own residual maps) can be found in Figure S8 and S9.

### Training on the un-residualised data is suboptimal to capture individual differences

Up to this point, we have shown that resting-state variations can fundamentally capture the inter-subject differences in task-evoked brain activity. The next question we asked is whether the model can recover individual idiosyncrasies in task-fMRI, if trained on the un-residualised resting-state spatial modes and the task activations, as opposed to the residualised data (i.e., variation maps). Having close-to-zero shared variance with the group-average, the residuals more accurately profile the individual differences by design; we posit that training on residuals avoids the contamination of group-level information and thus may potentially facilitate capturing individual-unique patterns. To fairly compare the two options requires recovering the actual task-evoked responses (as opposed to the residuals) from the predicted variations for each individual. To explore this, we next generated the surrogate activation amplitude using the PFMs’ amplitude for each individual, then added the group-average activation map (scaled by the resting-state-predicted amplitude) back to the predicted variation maps. These predictions with group-average added back in were correlated against the actual (un-residualised) activations for all subjects, again yielding a subject by subject correlation matrix per contrast. We calculated the prediction accuracy and discriminability from these correlation matrices, and compared them with the model trained on the un-residualised PFMs and task activations.

Overall, both options manifested considerable predictive power of individual activations, as suggested by the overall accuracy and discriminability (Figure 3, red and orange). Additionally, we found that although training on variations exhibited little improvement on the actual prediction accuracy (figure 3a and b), it tended to improve prediction discriminability (figure 3c and d). This suggests that it is more desirable to establish a mapping between the variations in rest and task data *per se* than simply use the original data with group-average effects present. This is probably because residualisation orthogonalises the individual maps with respect to the group-average maps and prevents the dominance of the typical activation patterns. Furthermore, this shows that separating out the modelling of overall amplitude from (group-mean-removed) map variability, and then recombining these parts of the model later, is at least as effective as predicting raw task from raw resting maps. This is valuable, as it does suggest that these different data aspects can indeed be considered separately. The subject identification accuracy based on residualised predictions (with group-average effects added back in evaludation) is shown in Figure S11 and Figure S12.

**Figure 3.**
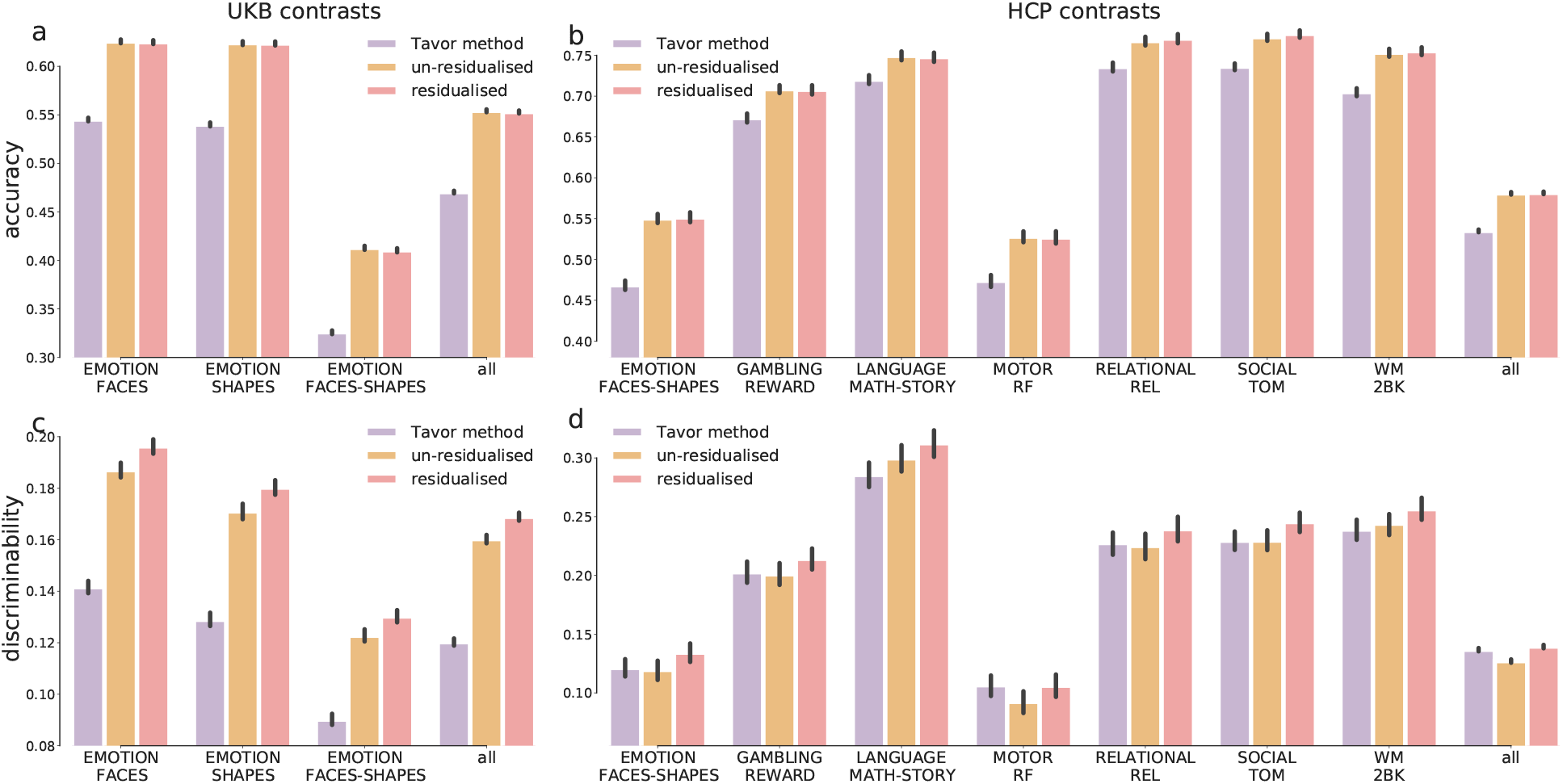
Comparison between the Tavor model and the ensemble models. Overall, the ensemble model trained on variation maps (residualised maps) outperformed the other two options; error bars show the 95% CI of the means. Prediction accuracy across 17,560 UKB subjects of the three contrasts, the last column showing all contrasts pooled together. (b) Equivalent plots across 991 HCP subjects of seven representative contrasts, the last column showing all 47 contrasts pooled together. (c) Prediction discriminability in UKB. (d) Prediction discriminability in HCP in a subset of task contrasts (see Figure S10 for all HCP contrasts).

We also benchmark our model against previous GLM-based methods (Tavor et al. 2016) using the same subjects. The Tavor method is based on multiple GLMs, essentially very similar to the baseline model, except for a few differences: (1) instead of training a global GLM for the whole brain between the resting-state and the task maps (as in our baseline model), the Tavor model seeks to fit multiple “local” GLMs within each of the pre-determined parcels; (2) the features of the Tavor model are seed-based connectivity maps, while our baseline model uses the dual-regression maps (i.e., multiple regression against the many “seed” timeseries output by the first stage of dual-regression). The ensemble model, trained either on the un-residualised data or on the variation maps, yielded higher prediction accuracy than the Tavor method. On the UKB dataset, the ensemble model substantially improved prediction accuracy and discriminability; on the HCP dataset, the Tavor method and the ensemble model trained on variations manifested comparable discriminability, both superior to the ensemble model trained on un-residualised data (see Figure S10 for all HCP task contrasts). Note that, among the HCP contrasts, motor-tasks exhibited weak prediction discriminability.A possible explanation for this is that the individual response profiles to motor-related stimulus had little cross-subject variations, such that the model was not able to extract sufficient information to discriminate between subjects. The relatively lower prediction accuracy of motor tasks is, on the other hand, unexpected, especially considering the strong activations in cortical regions that are supposed to enable the model to learn the mapping between resting-state and motor tasks. Understanding this discrepancy between motor tasks and resting-state activity requires future investigations and would be important to understand the ongoing interplay of resting-state networks in task execution.

The fact that the model trained on the variations *per se* (with an explicit and separate amplitude prediction) can better capture patterns unique to individuals than its un-residualised counterpart corroborates the assumption that, in addition to the spatial layout (shapes) of activations, the overall activation intensity also contributes to the variability of task-elicited activity. Following this, we also tested whether incorporating resting-state amplitude as additional predictors explicitly at the ensemble stage would further facilitate capturing individual-unique patterns for the un-residualised model. We found that, though having little effect on the actual prediction accuracy, including the PFMs’ amplitude as explicit predictors (in addition to the other two predictors, the baseline-model predicted and sparse-model-predicted values in the corresponding voxel) did further improve discriminability (Figure S13a and b). This again supports our findings that the inherent variations in resting-state and task activity are more informative of the mapping between the two states than the original activity profiles. For the ensemble model trained on the residualised data, regressing out the group-average response “removes” the overall activation intensity relative to the group-average activations for each individual. Therefore, introducing resting-state amplitude to the residualised ensemble model, in theory, should have little effect on model performance. However, in practice, we found that incorporating resting-state amplitude as additional features in the ensemble stage also increased prediction discriminability for the residualised ensemble model. There are a few possible explanations for this discrepancy. One possible explanation is that the group-average activation patterns were not entirely removed particularly from the subjects that are very atypical, probably due to GLM’s sensitivity to outliers or noise in the fitting (e.g., related to regression dilution). In this sense, including resting-state amplitude as additional features thus accounted for the remnants of the amplitude information particularly for those atypical subjects, and thus increased the overall prediction discriminability (Figure S13d and f) on the UKB dataset. Another possibility is that the overall activation intensity may still inform the (strength of the) variabilities of the shape of activations. This possibility can be partially validated by the findings that it further improved the fit with the spatial pattern of inter-individual variability by including resting-state amplitude as additional features at the ensemble stage (Figure S13c and f). Note that, however, the resting-state amplitude is not expected to be a perfect surrogate of the task amplitude. The *R*^2^ between the actual and the predicted task amplitude is actually small (Figure S1 and S2).

### Prediction accuracy paralleled test-retest reliability

To evaluate whether the predicted task maps can reliably capturing individual differences in tasks, we utilised retest scans in HCP data to compare the prediction accuracy of task maps against test-retest correlations of tasks. The second-scan task contrast maps (either residualised or un-residualised) were correlated against the first-scan task maps for subjects that had received second-time scanning, yielding a subject by subject correlation matrix per given contrast. We also investigated the reliability of activation amplitude by residualising the second-time task contrast maps using the original (time 1) group-average map, and correlated the amplitude values (i.e., regression betas) against the first time task amplitude. We tested whether resting-state-predicted amplitude is more robust than those measured directly in tfMRI.

For both datasets, the PFM-predicted contrast maps yielded higher overall accuracy than the repeat scans, consistent across all task contrasts (Figure 4a and c, light blue and light white), suggesting that resting-state predicted activations can surpass task-fMRI retest reliability. This coincides with previous studies that resting-state features serves as a reliable trait marker and may even be more heritable than task-fMRI phenotypes (Winkler et al. 2010). Note that, the accuracy of PFM-predicted activations that is on par with the test-retest reliability is unlikely a result of over-fitting to the first-visit tfMRI data. With the repeat scans entirely invisible to training, the PFM-predicted task activations still generalised well to the second-visit task contrast maps (see Figure 4, light green bars); actually, the PFM-predicted task maps (predicted using the first visit resting maps only) gave comparable prediction accuracy for both visits. Furthermore, the PFM-predicted task amplitude proved more reliable to task-fMRI scans in replicating the overall activation amplitude (Figure 4b and d).

**Figure 4.**
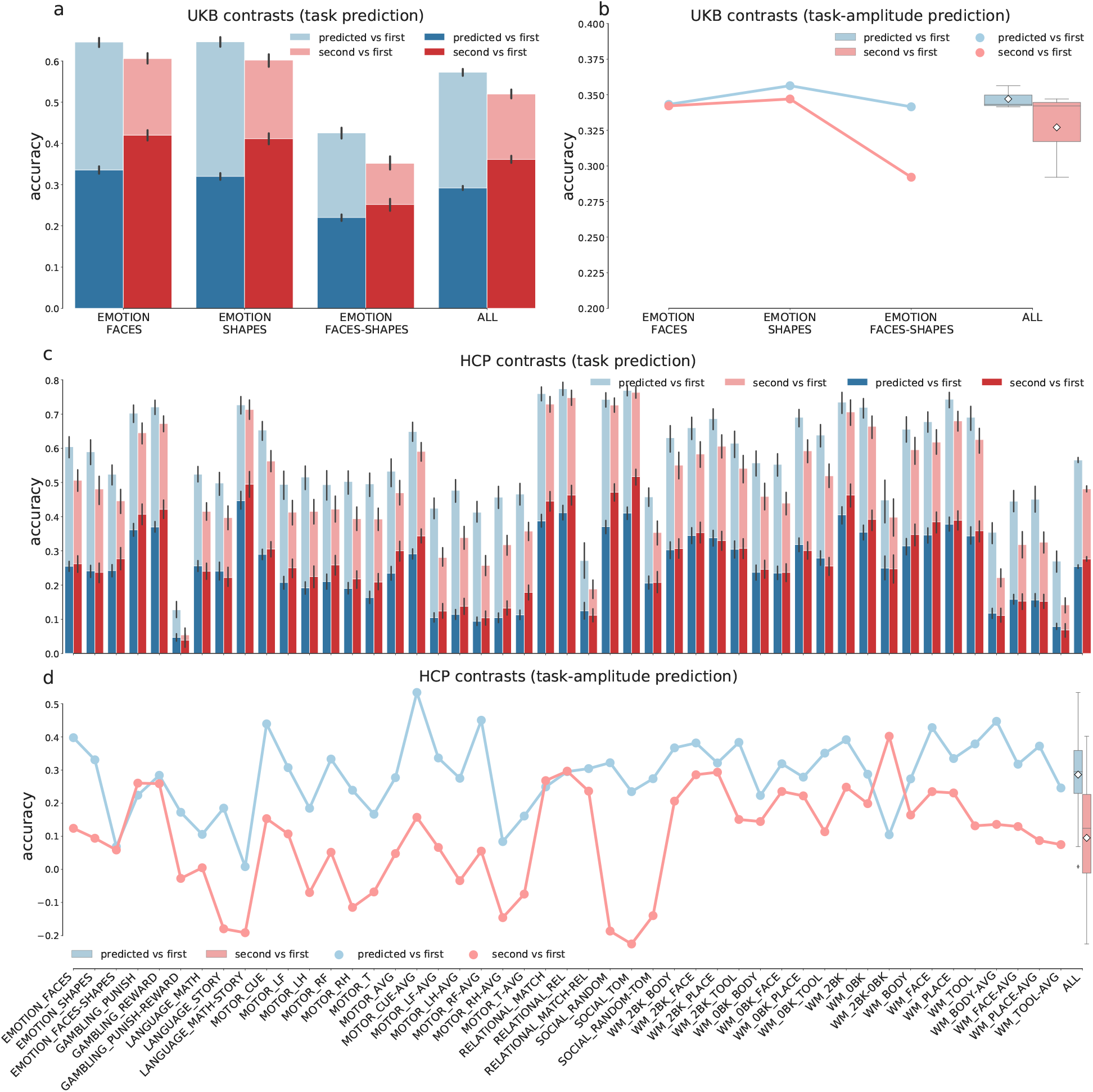
Test-retest reliability of PFM-predicted task maps. In (a) and (c), dark colors denote accuracy of residual predictions; pale colors the accuracy of group-average-added-back predictions. Blue: accuracy of PFM-predicted maps. Red: accuracy of the second-visit tfMRI contrast maps. Although the group-average-added-back predictions consistently yielded higher accuracy than the retest scans, on UKB the accuracy of residual predictions is yet to be improved. On HCP, in contrast, the accuracy of residual predictions was approaching the second-visit scans, possibly due to the much longer scanning sessions. (b) and (d). For both datasets, PFM-predicted task amplitude was overall more reliable than the second-time task-fMRI scans.

As mentioned in previous sections, predicting residual variation is of more interest. On the HCP dataset, the accuracy of residualised predictions approaches the test-retest reliability of task variation maps (Figure 4c) for most contrasts, and yielded higher accuracy for several contrasts (GAMBLING_REWARD, GAMBLING_PUNISHMENT, SOCIAL_MATCH-REL, etc.). On the UKB dataset, however, the re-test (residualised) tfMRI scans still yielded much higher accuracy than the PFM-predicted task variations (Figure 4a), possibly because of the much shorter resting-state scanning sessions. The retest scans also had higher prediction discriminability than did the group-average-added-back predictions, which is un-surprising due to the dominance of group-average effects (Figure S16).

Figures 5 and 6 show the comparison between the predicted, actual, and group-average activations volumetrically (for UKB) and on the surface (for HCP). It can be seen that the predicted activations provide a “smoothed” estimation of the individual activations, while preserving the unique patterns in individual subjects (for the actual and the predicted task variation maps of the same example subjects, see Figure S17 and S18).

**Figure 5.**
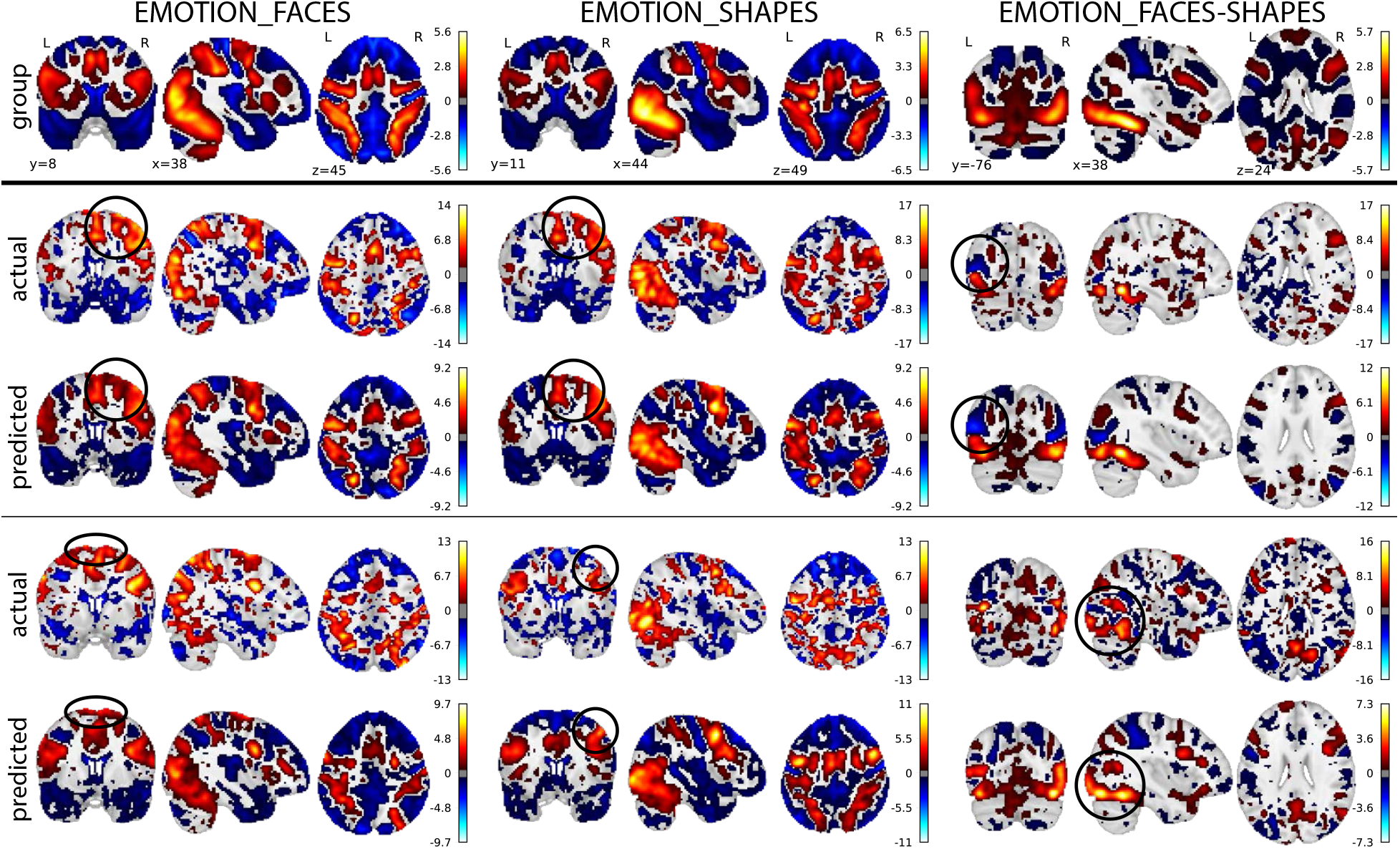
Predicted, actual, and group-average activations of 6 example UKB subjects. The predicted activations captured the atypical activations in individual subjects (with group-mean-related components included). The subjects ranked between 50% to 75% according to their correlations with the corresponding group-average activations. See Figure S17 for the plots of the predicted and the actual task variation maps of the same example subjects.

**Figure 6.**
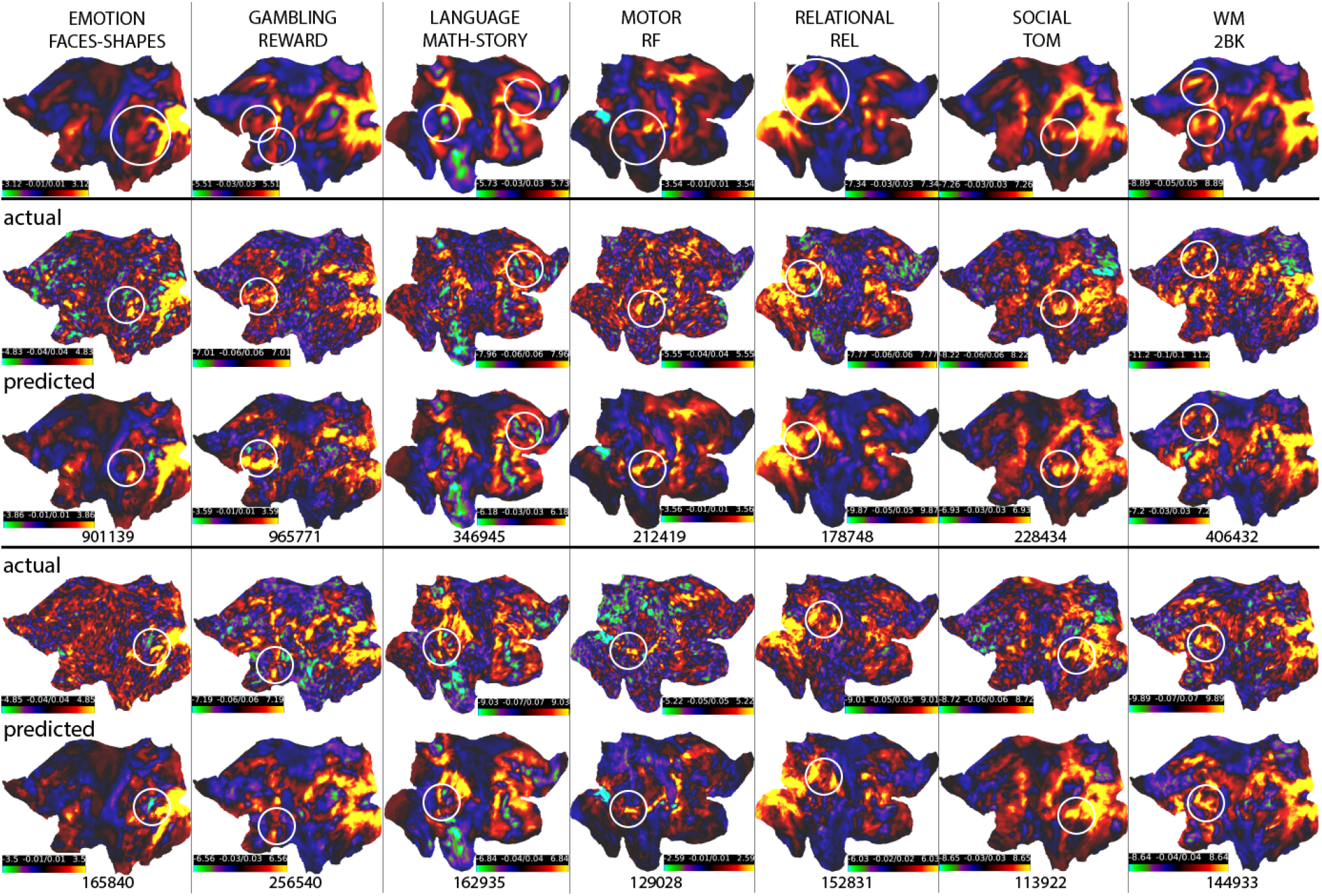
Predicted, actual, and group-average activations of example HCP subjects. The predicted activations captured the atypical activations in individual subjects; these subjects ranked in the lower 50% percentile according to their correlations with the corresponding group-average activations. See Figure S18 for plots of the actual and the predicted task variations maps of the same example subjects.

## DISCUSSION

In this paper, we extended previous GLM-based approaches (Tavor et al. 2016; Cohen et al. 2020; Dohmatob et al. 2021) and proposed an ensemble learner to model individual variations in task activations on two large datasets, UKB and HCP. Enabled by a recently developed technique, sPROFUMO, we exploited the richness of individual variability in resting-state to reproduce task-evoked activation patterns unique to individuals. We demonstrated that sPROFUMO can accommodate higher predictive power than DR-ICA, especially in terms of the overall capacity of reproducing between-subject differences. This added advantage of sPROFUMO arises from its enhanced ability to depict fine-grained resting-state variability in rich detail due to its bidirectional and hierarchical architecture between the group-average and individual,in contrast to the unidirectional group-average algorithms (e.g., DR-ICA). Furthermore, we showed that modelling the individual activation profiles as a combination of the group-average and predicted variations can be more desirable than simply modelling the raw task map, suggesting that two sources of task variability, shape and amplitude, factorise into different compartments and can be modelled separately. Characterising different aspects of task variability is important to understanding the sources of these cross-subject differences. Overall, resting-state functional modes serve as a set of bases that can not only sufficiently reconstruct individual task-fMRI space but also yield more reliable localisation of individual task-evoked response profiles.

Our ensemble framework consists a baseline model and a sparse model, each tailored to a different scenario. In the baseline model, for each individual, the resting-state modes span the space of the task activation maps and thus, in theory, can reproduce task-fMRI in itself. In practice, however, more spatial complexity is often required to select local features that are “cleaner” or of more interest. The sparse model largely accounts for this limitation. For example, the motor network in resting-state modes contains components that are often in sync with each other and are part of the same spatial basis. The baseline model cannot split them, while the sparse model may select the components more desirable for prediction. However, the sparse model has another caveat. Despite the existing rescaling techniques (e.g., fitting another OLS on top of the selected features; introducing a re-scale factor), the Lasso penalty often introduces too much shrinkage, particularly when the prediction involves too many candidate features. As a results, the predicted response may become too biased towards zero thus degrading model discriminability. The ensemble model, by fitting another OLS on top of each voxel, de-biases the over-shrinkage of the sparse model.

It is worth noting that the group-average activation patterns alone can have considerable overlap with individual activation maps. Thus one can obtain moderate prediction accuracy by simply reproducing the group-average. Hence, the accuracy of residualised predictions, or the discriminability of the group-average-added-back predictions, are more informative on the model’s ability to make individualised predictions. This is of particular importance, because many existing algorithms tend to push predictions towards the mean. In a higher-dimensional setting, the relation between the two measures becomes complicated, but it is not difficult to see that the improvement of discriminability may degrade accuracy a little. Training and evaluating the model on residualised resting-state and task data thus have more desirable properties, not only to simplify the assessment of model performance but also to maximise ability to capture inter-individual differences. Other approaches to improve prediction discriminability include introducing a contrastive loss term to push between-subject differences to be large (Ngo et al. 2021). It is yet to be investigated whether the two approaches are comparable. However, introducing extra terms may complicate the loss function (for example, turn a convex loss function into a non-convex one) and thus may be less stable. Training on residualised data keeps the original loss function structure and is usually simplier to train.

In addition to predicting individual-unique activations, it is also of value to investigate the causes of the variations in task-evoked activations, particularly, what information in resting-state activity drives the individual differences in task activity. For example, do variations in peak activation patterns correspond to the changes in resting-state activity in the same location, or is it actually driven by more complicated configuration of the dense connectivity pattern? Such investigations would help us understand the nature of the inherent resting-state features that characterise variations in task activity. For example, these features can be “structurally” inherent (characterised by brain organization and connectivity) or “functionally inherent” (related to the cognitive state of subjects during the resting-state scan) (Tavor et al. 2016), both of which may cause the re-configuration or reallocation of peak activation patterns. Note that, individual differences in task-evoked activations may be partially due to inter-subject misalignment. Indeed, registration remains an empirical question and may be sub-optimal in practice. However, it is very unlikely that our results only account for misalignment between subjects, as the model can capture variations not only in shape and position but also in topology of the activations. Indeed, it is likely that the relatively state-of-the-art alignments used here in preprocessing reduced intersubject variability, rather than increased it.

Using resting-state fMRI scans to infer individualised task-evoked response has a wide range of implications in translational and clinical neuroscience. One potential application of the proposed model is to infer individualised functional localisers based on resting-state fMRI scans. This is important because tfMRI scans are often of limited accuracy and reliability (Elliott et al. 2020; Ellis et al. 2020), possibly due to poor task performance and noise that is hard to remove in pre-processing. Such a framework can supplement task localisers, potentially improving the prediction of individual functional mapping and facilitating investigations of individualised response profiles of localised brain regions. Furthermore, as numerous multi-site multi-scanner consortia emerge, it is also important to reduce scanner-induced or age-induced bias such that the model can be generalised beyond sites or populations. This requires efforts to develop a model that is capable of learning features invariant across scanners and insensitive to confounds. If generalisable to other populations, such a model can be used to localise regions of interest for those who cannot perform tasks, such as paralysed patients and infants.

There are a few limitations in this study. First, the ensemble model is a linear combination of two single (largely) linear models and thus has limited ability to capture higher-order non-linear relationships between the resting-state and task-evoked brain activity. Second, the decompositions of common modes of variations are unsupervised. In the future, more complex modelling could be adopted to simultaneously estimate the common modes of variations and the reconstruction coefficients. Third, the rich information derivable from T1 and diffusion MRI scans may further aid the predictions of individual differences in task-evoked activity, and this model is yet to be adapted into a multi-modal framework.

## CODE AVAILABILITY

Code for the model and analysis in this paper can be found in https://github.com/yingqiuz/predict-task-individual-variability.

Code for obtaining PFMs will be made available in an upcoming FSL release. It is currently available in https://git.fmrib.ox.ac.uk/rezvanh/sprofumo_develop.

## ACKNOWLEDGEMENTS

SMS is supported by the Wellcome Trust Collaborative Award (215573/Z/19/Z), and MRC Mental Health Pathfinder grant (MC_PC_17215). SJ is supported by a Wellcome Senior Fellowship (221933/Z/20/Z) and Wellcome Collaborative Award (215573/Z/19/Z). The Wellcome Centre for Integrative Neuroimaging is supported by core funding from the Wellcome Trust (203139/Z/16/Z). The computational aspects of this research were partly carried out at Oxford Biomedical Research Computing (BMRC), that is funded by the NIHR Oxford BRC with additional support from the Wellcome Trust Core Award Grant Number 203141/Z/16/Z. We are grateful to UK Biobank for making this invaluable resource available (data access application 8107), and to the UK Biobank participants for dedicating their time to make this data possible. We are also grateful to the Human Connectome Project, WU-Minn Consortium (Principal Investigators: David Van Essen and Kamil Ugurbil; 1U54MH091657) funded by the 16 NIH Institutes and Centers that support the NIH Blueprint for Neuroscience Research; and by the McDonnell Center for Systems Neuroscience at Washington University. We additionally thank Mark Chiew and Diego Vidaurre for their helpful discussions.

## SUPPLEMENTARY INFORMATION

**TABLE S1.** List of the 47 HCP contrasts. We used the 47 unique contrast maps for HCP, excluding all redundant contrasts.

| HCP contrast category | HCP contrast index and name |
| --- | --- |
| EMOTION | 01_EMOTION_FACES 02_EMOTION_SHAPES<br>03_EMOTION_FACES-SHAPES |
| GAMBLING | 07_GAMBLING_PUNISH 08_GAMBLING_REWARD<br>09_GAMBLING_PUNISH-REWARD |
| LANGUAGE | 13_LANGUAGE_MATH 14_LANGUAGE_STORY<br>15_LANGUAGE_MATH-STORY |
| MOTION | 19_MOTOR_CUE 20_MOTOR_LF 21_MOTOR_LH 22_MOTOR_RF<br>23_MOTOR_RH 24_MOTOR_T 25_MOTOR_AVG<br>26_MOTOR_CUE-AVG 27_MOTOR_LF-AVG 28_MOTOR_LH-AVG<br>29_MOTOR_RF-AVG 30_MOTOR_RH-AVG 31_MOTOR_T-AVG |
| RELATIONAL | 45_RELATIONAL_MATCH 46_RELATIONAL_REL<br>47_RELATIONAL_MATCH-REL |
| SOCIAL | 51_SOCIAL_RANDOM 52_SOCIAL_TOM<br>53_SOCIAL_RANDOM-TOM |
| WORKING MEMORY | 57_WM_2BK 58_WM_2BK 59_WM_2BK 60_WM_2BK<br>61_WM_0BK 62_WM_0BK 63_WM_0BK 64_WM_0BK<br>65_WM_2BK 66_WM_0BK 67_WM_2BK-0BK 71_WM_BODY<br>72_WM_FACE 73_WM_PLACE 74_WM_TOOL 75_WM_BODY-AVG<br>76_WM_FACE-AVG 77_WM_PLACE-AVG 78_WM_TOOL-AVG |

**TABLE S2.**
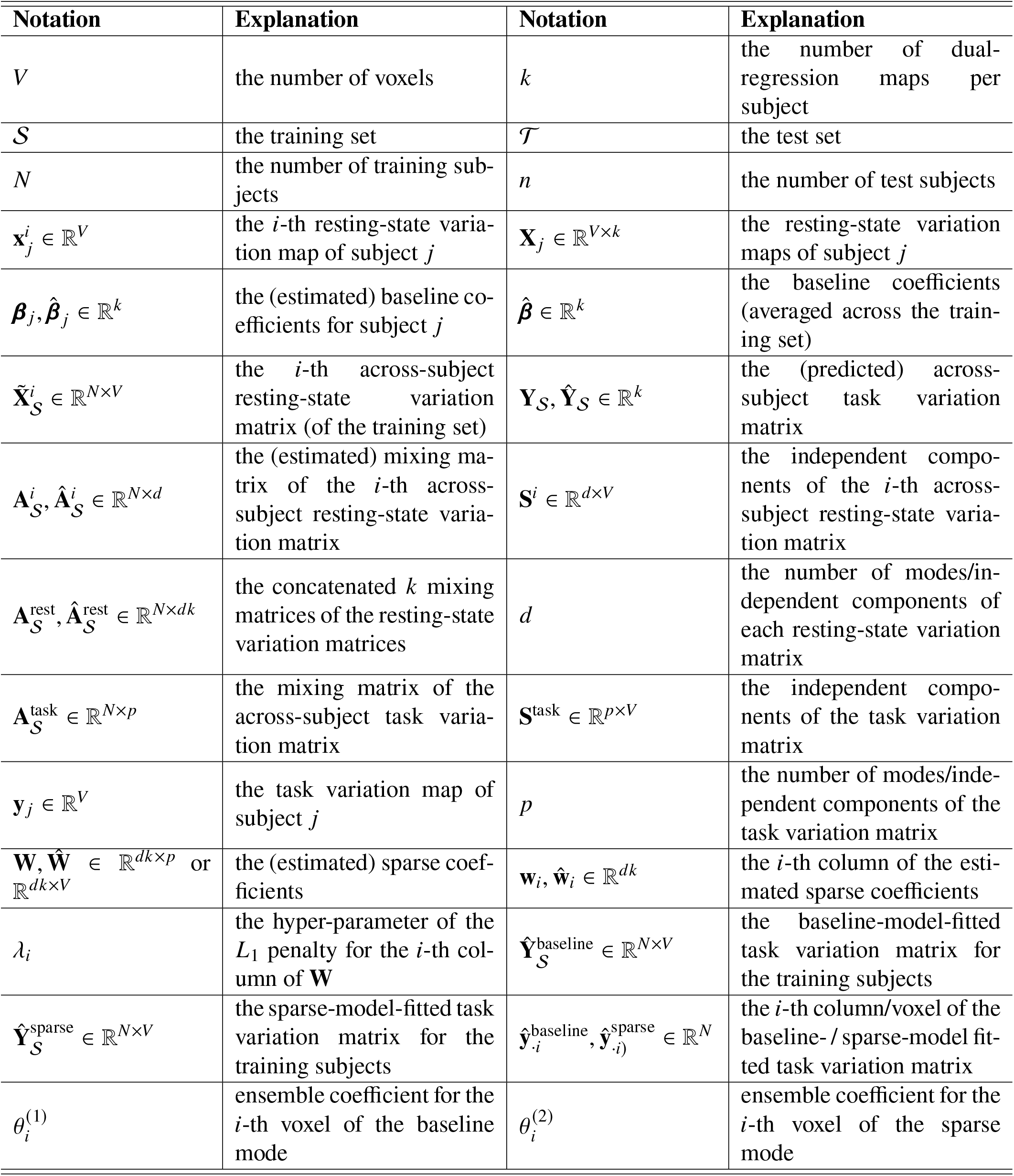
List of the notations.

**Figure S1.**
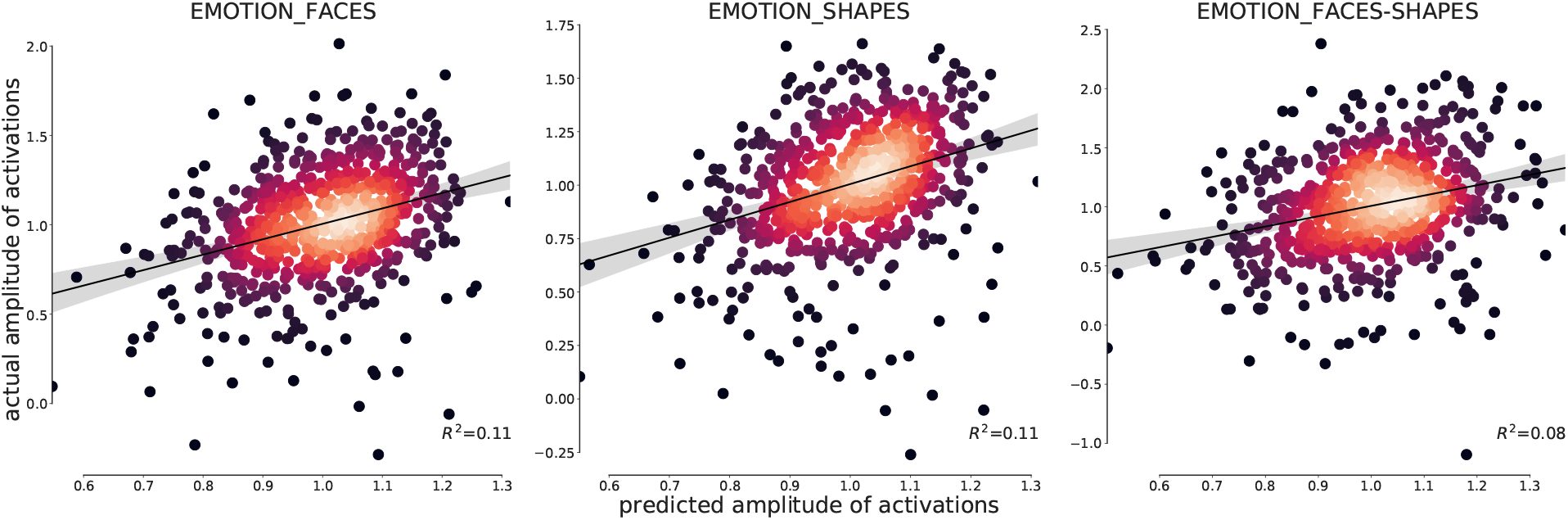
Using resting-state amplitude to predict activation amplitude (UKB). For each task contrast, the activation amplitude was predicted using the amplitude of the 50 PFMs (700 subjects shown).

**Figure S2.**
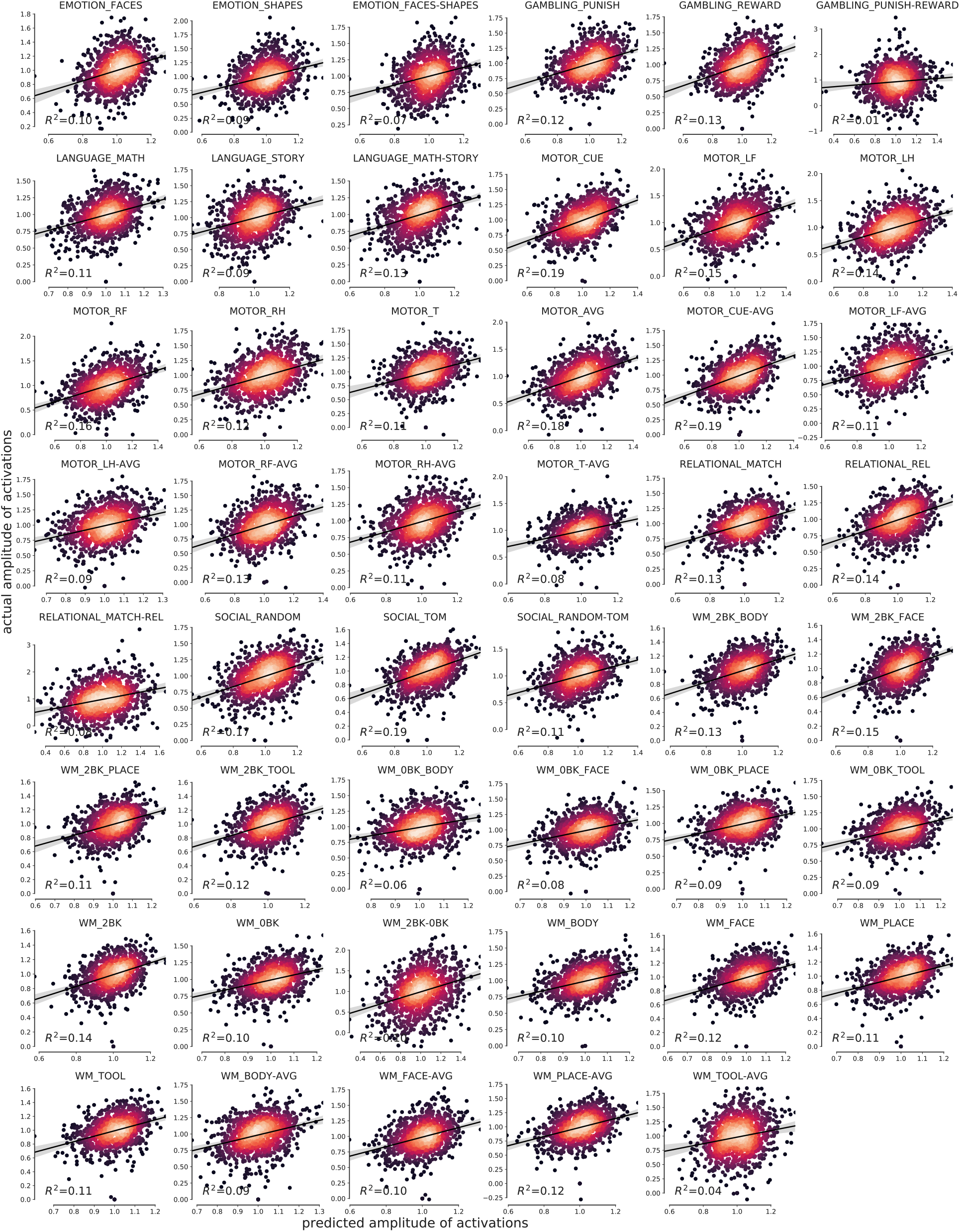
Using resting-state amplitude to predict activation amplitude (HCP). For each task contrast, task amplitude was predicted using the amplitude of 150 PFMs via 10-fold cross-validation (i.e., trained on 9 folds and predicted on the rest).

**Figure S3.**
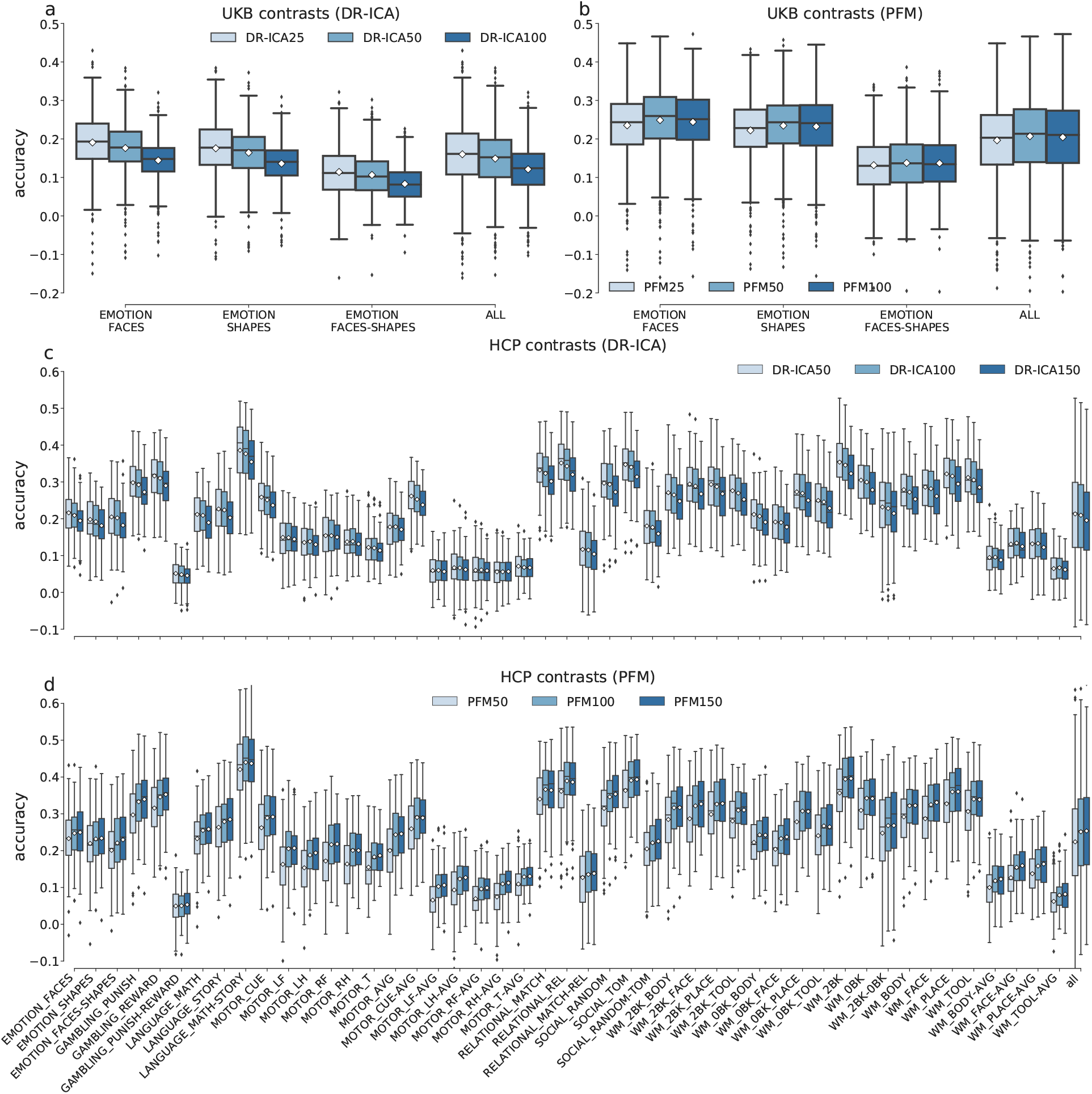
Choices of the functional modes’ dimensions. Lower dimensions often result in larger parcels and tend to reflect whole brain resting-state networks. Higher dimensions, in contrast, tend to break down the parcels into more fine-grained functional subprocesses. We argue that the resolution of functional parcellations may non-trivially impact prediction of individual variations in task-evoked activations. The number of resting-state modes must be optimised in the first place for the the subsequent analysis. White diamond shows the mean. (a) Prediction accuracy of DR-ICA25, DR-ICA50, and DR-ICA100 across a subset of 700 UKB subjects. (b) Equivalent plots of PFM25, PFM50, and PFM100. (c) and (d) Equivalent plots of 98 HCP subjects at 50, 100, and 150 modes. The results were based on the residualised data; for the un-residualised data, different dimensions had similar effects on the accuracy, though with smaller differences (not shown here).

**Figure S4.**
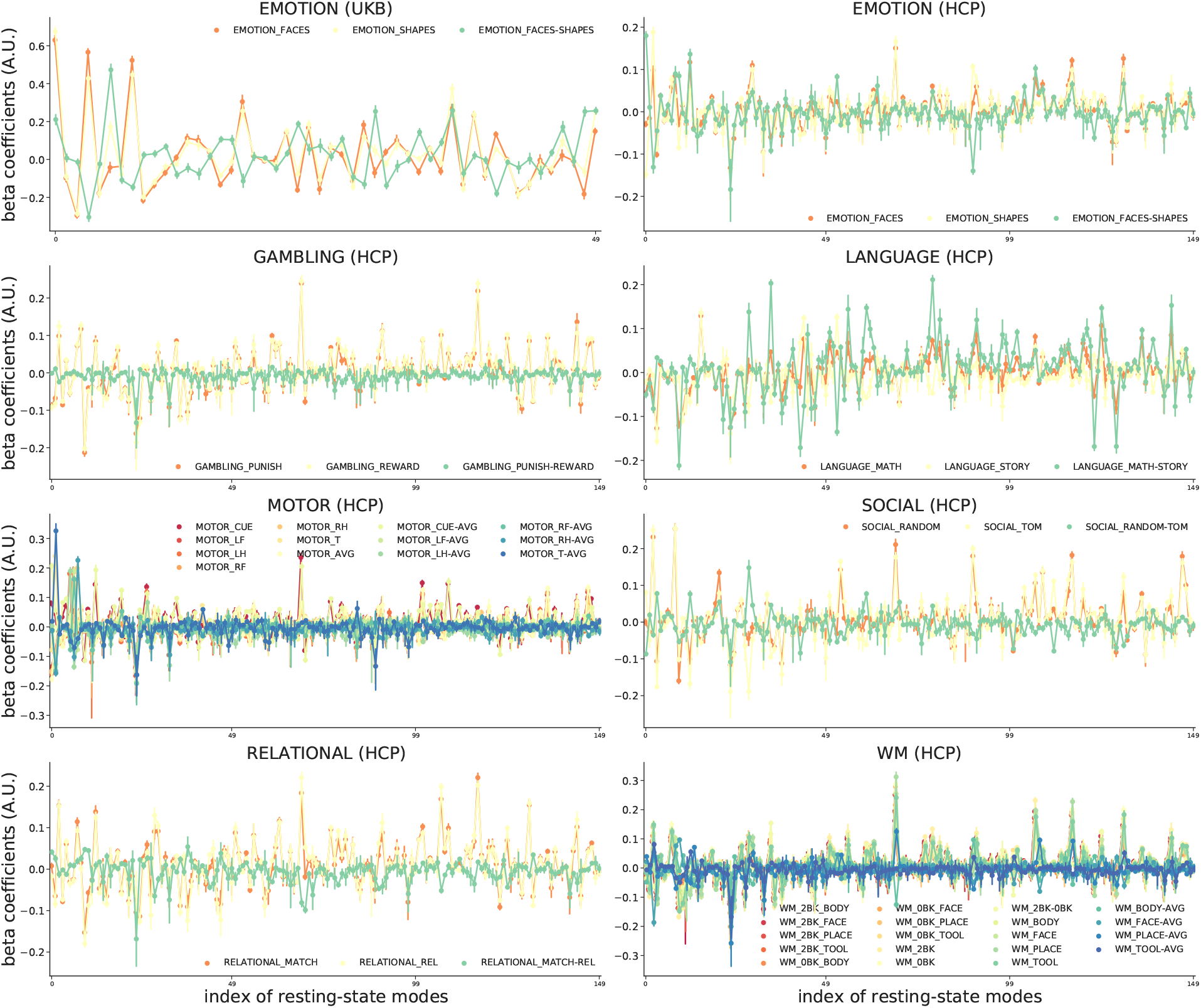
Baseline coefficients (betas of the baseline model) of the residualised data. Error bars showing 95% CI of the mean beta values (calculated across 1,000 UKB subjects and 891 HCP subjects). For each subject, the coefficients were divided by the maximum beta value within the given contrast. Overall, most functional modes exhibited consistent patterns within each task domain.

**Figure S5.**
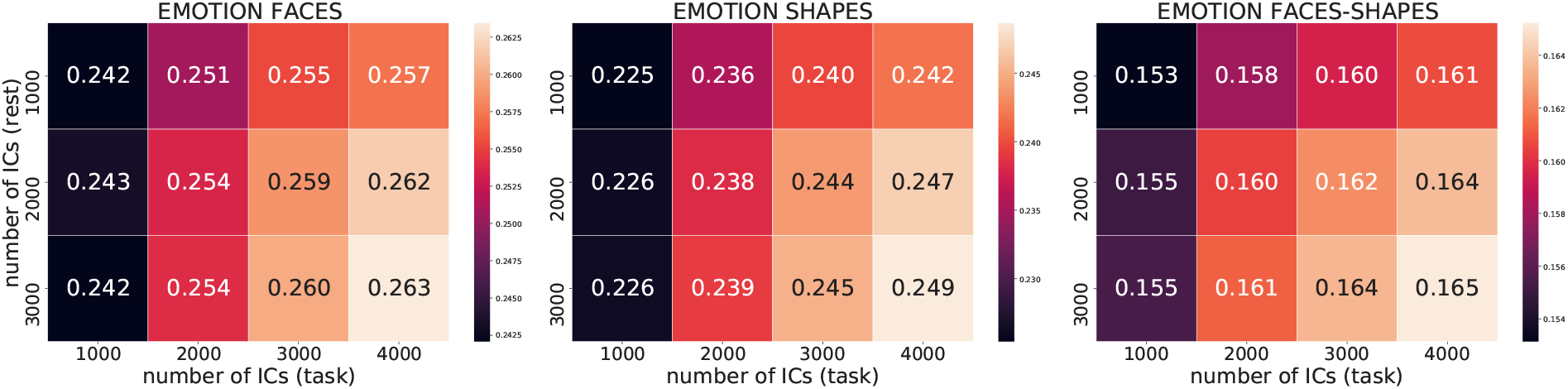
Prediction accuracy of the sparse model at a range of PFM dimensions, trained on a subset of 4,000 UKB subjects and tested on 700. Overall, prediction accuracy increases with the number of functional modes. Note that the results were based on residualised data. The un-residualised data exhibited similar accuracy patterns, though with smaller differences between the choices of dimensions (not shown here).

**Figure S6.**
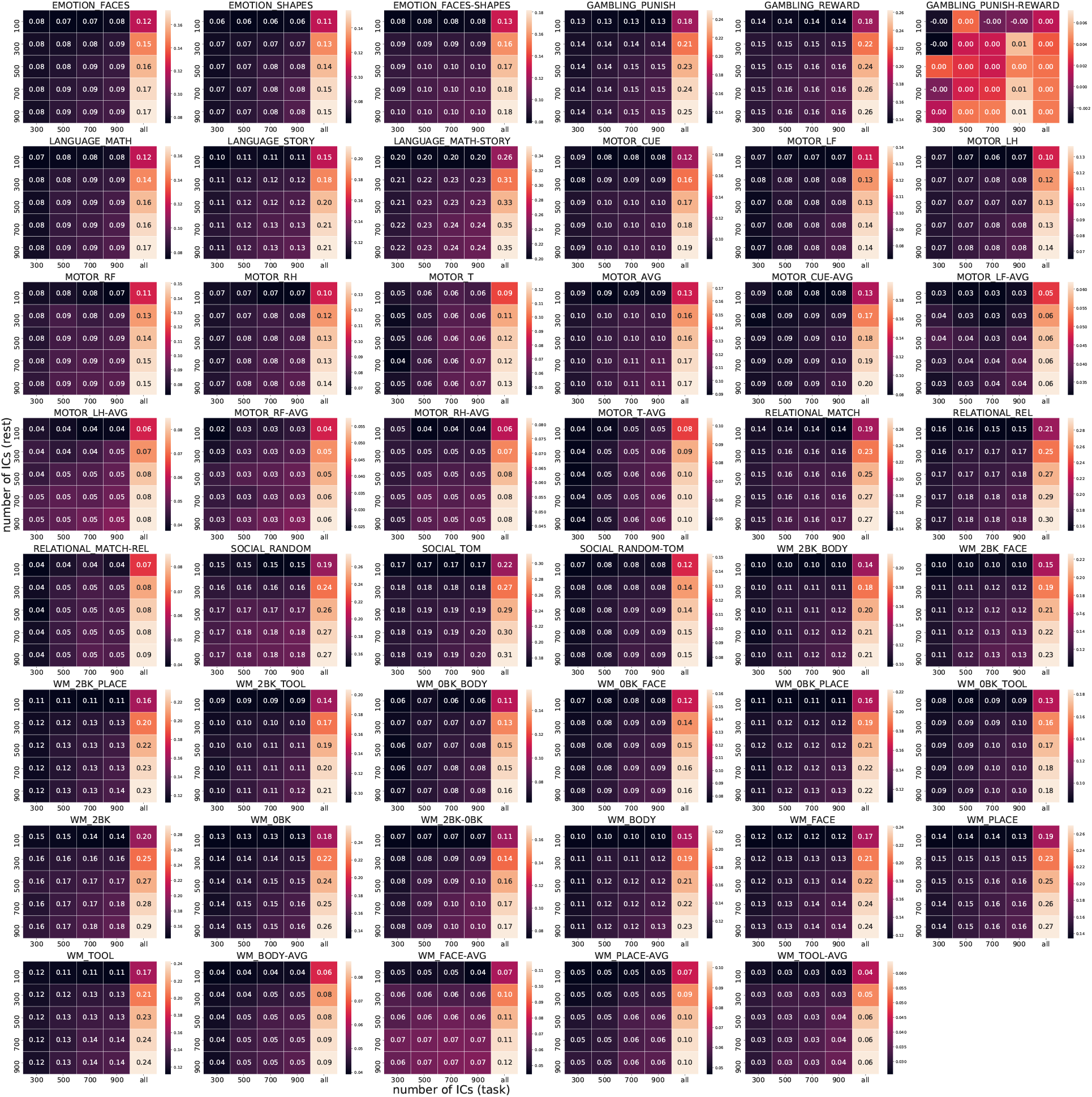
Prediction accuracy of the sparse model at a range of PFM dimensions, trained on 891 HCP subjects and tested on 98. Overall, accuracy increases with the number of functional modes. The un-residualised data exhibited similar accuracy patterns, though with smaller differences between the choices of dimensions (not shown here).

**Figure S7.**
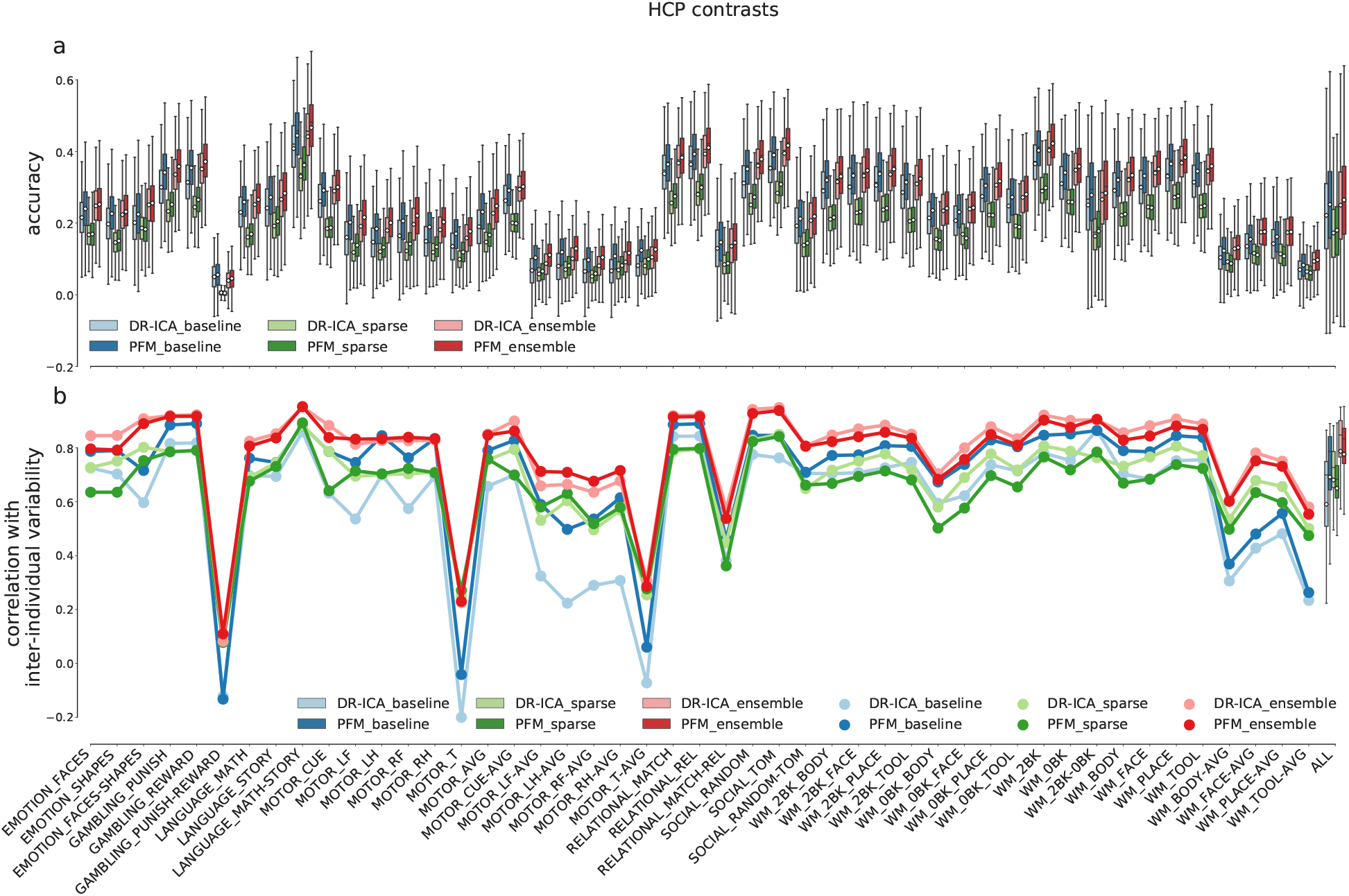
Comparison between the ensemble model and the single models, shown across all 47 HCP task contrasts. (a) Equivalent plots of Figure 2b. (b) Equivalent plots of Figure 2d.

**Figure S8.**
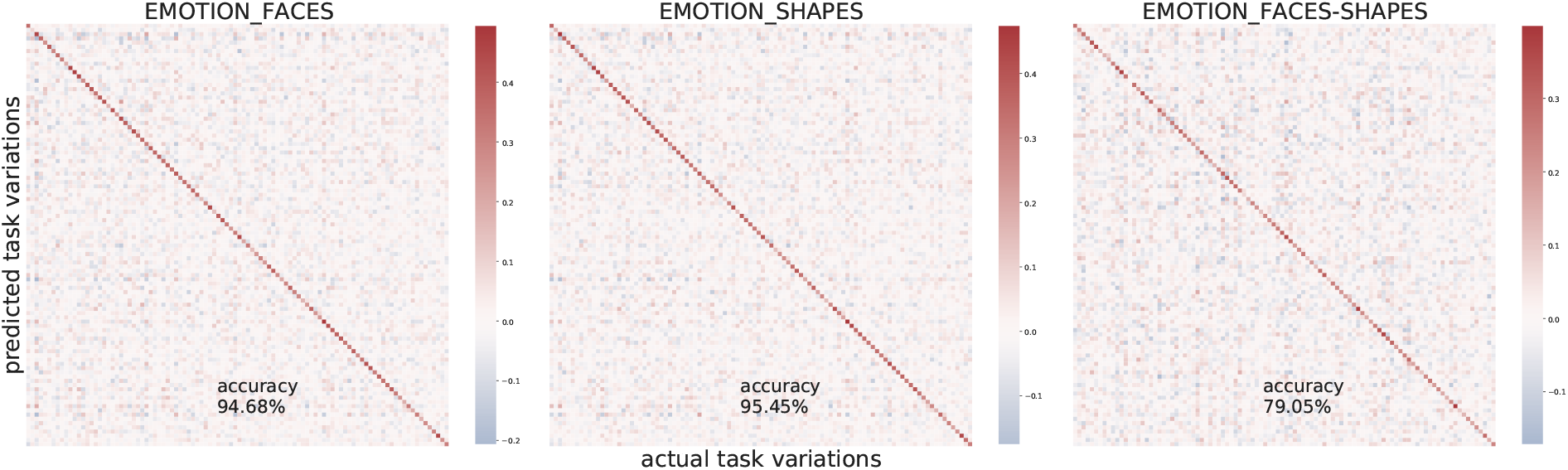
UKB subjects identification accuracy (based on residualised data). Accuracy (pearson’s correlation between the predicted and actual variations) was calculated across all subjects; for illustration purpose, only 100 subjects were shown above. The off-diagonal elements fluctuate around zero, i.e., accuracy and discriminability calculated on residualised predictions are almost identical.

**Figure S9.**
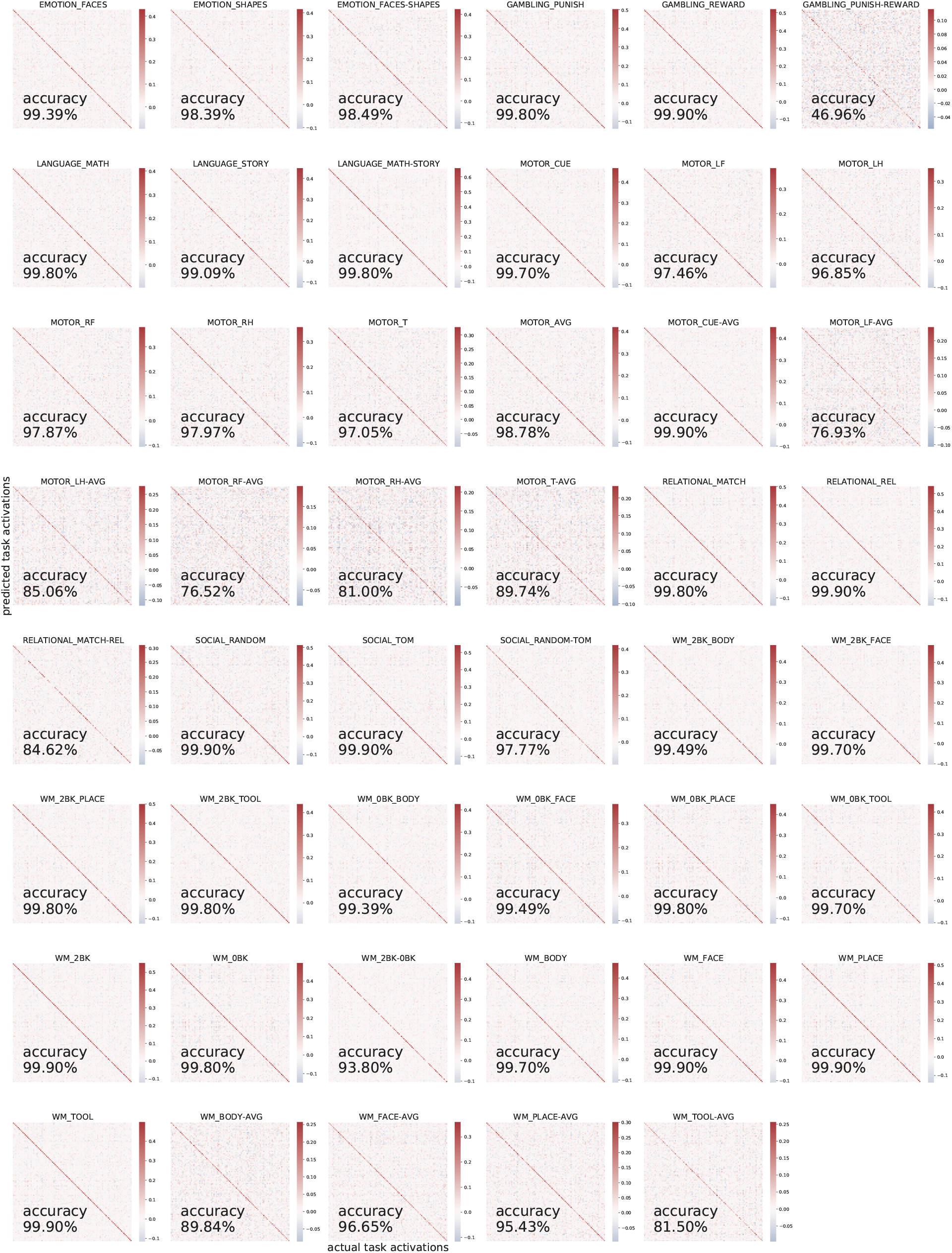
HCP subjects identification accuracy (based on residualised data). For illustration purpose, only 100 subjects were shown above.

**Figure S10.**
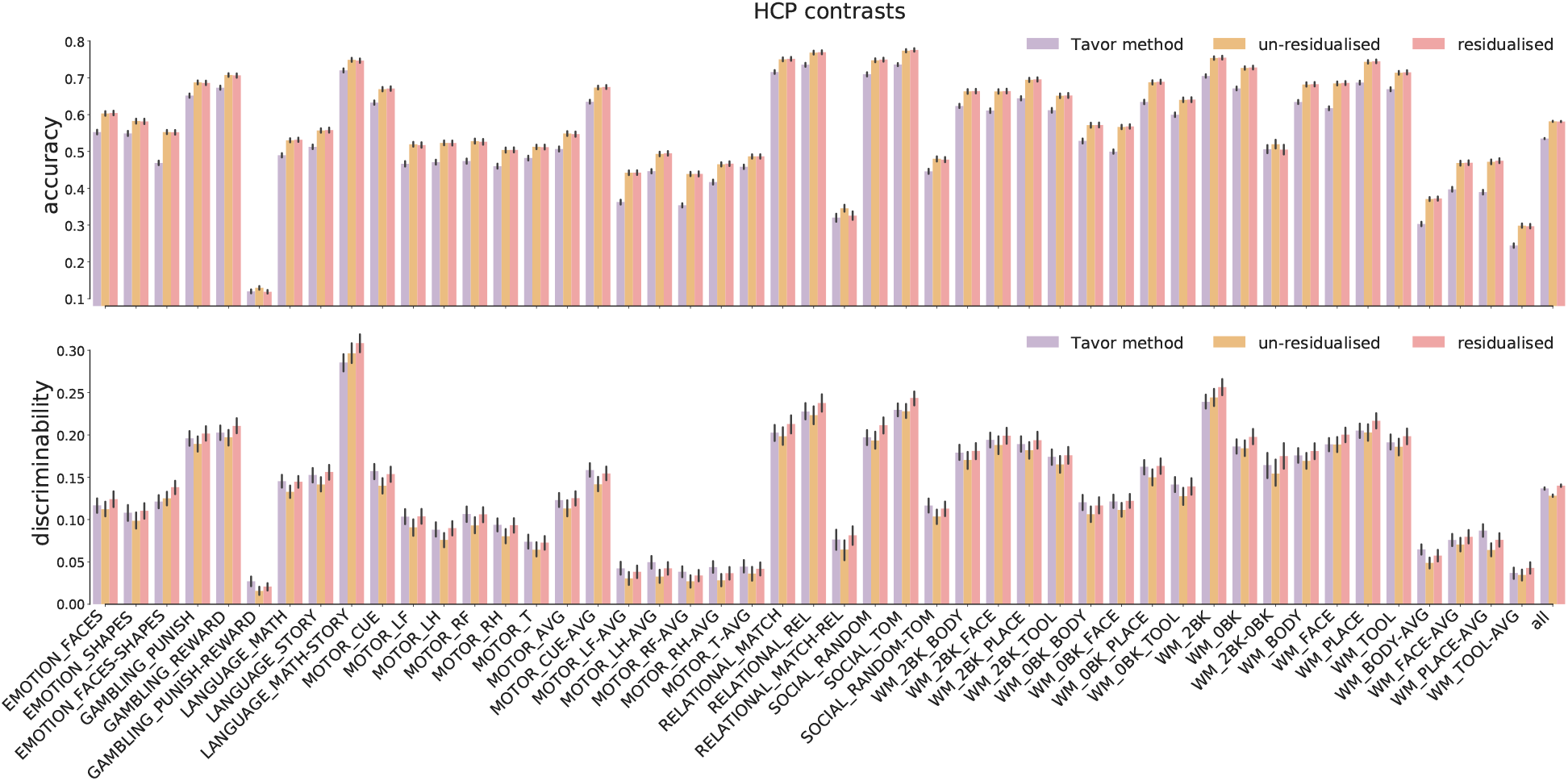
Comparison of the Tavor model and the ensemble model (both un-residualised and residualised) across 991 HCP subjects for all 47 task contrasts. The ensemble model (either residualised or not) outperformed the Tavor model in terms of the actual prediction accuracy; however, the Tavor model could make more individualised predictions than the ensemble model if both trained on un-residualised data. The residualised ensemble model outperformed the other two both in accuracy and discriminability, except for the motor task domain.

**Figure S11.**
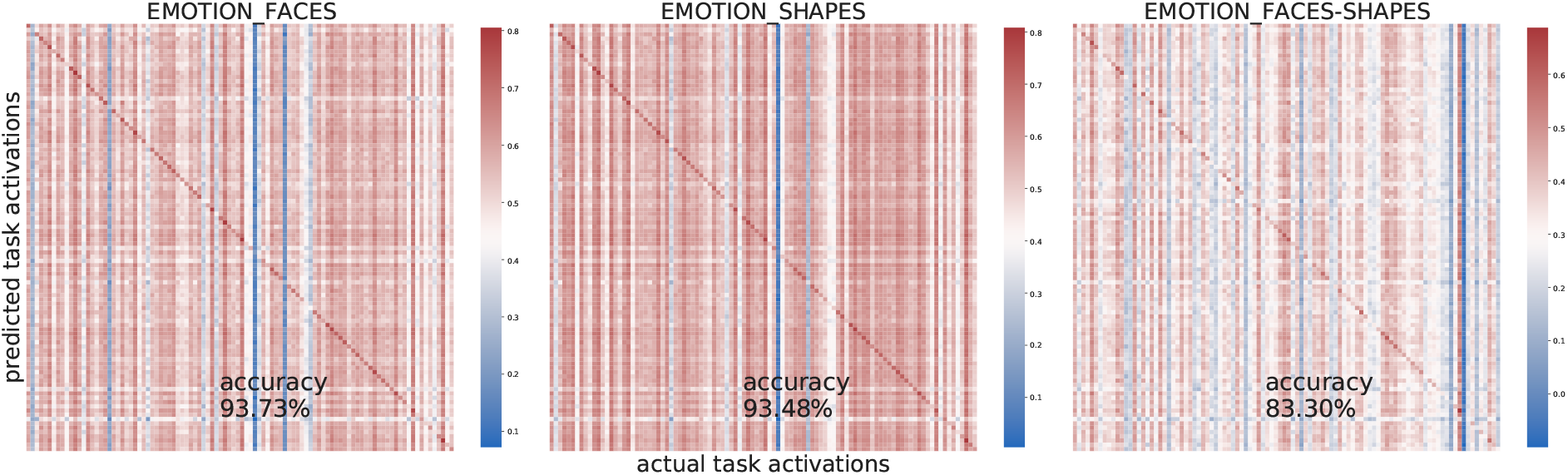
UKB subjects identification accuracy (with group-average activations added back in). The off-diagonal values no longer fluctuate around zero. The subject identification accuracy remains high. For illustration purpose, only 100 subjects were shown above.

**Figure S12.**
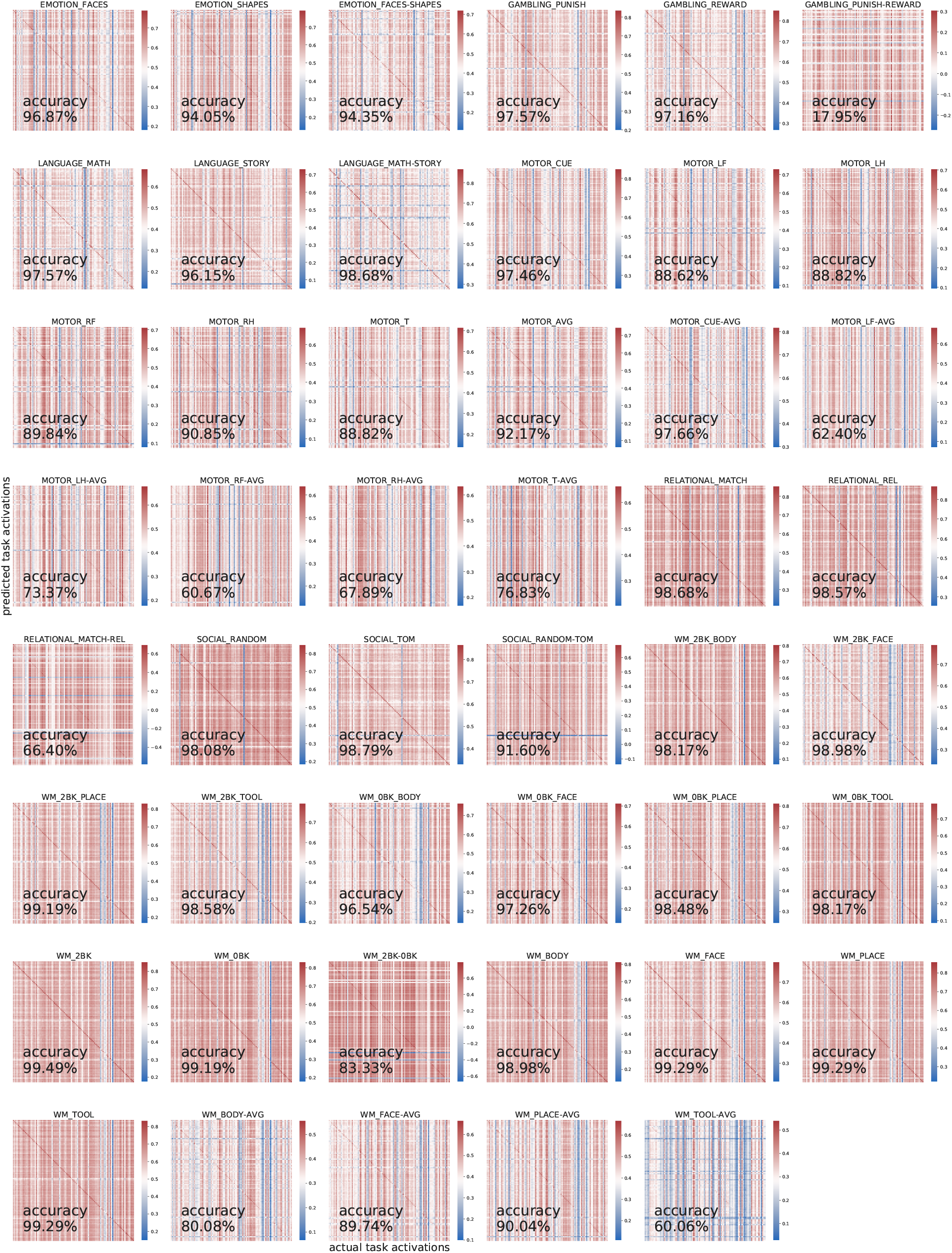
HCP subjects identification accuracy (with group-average activations added back in). For illustration purpose, only 100 subjects were shown above.

**Figure S13.**
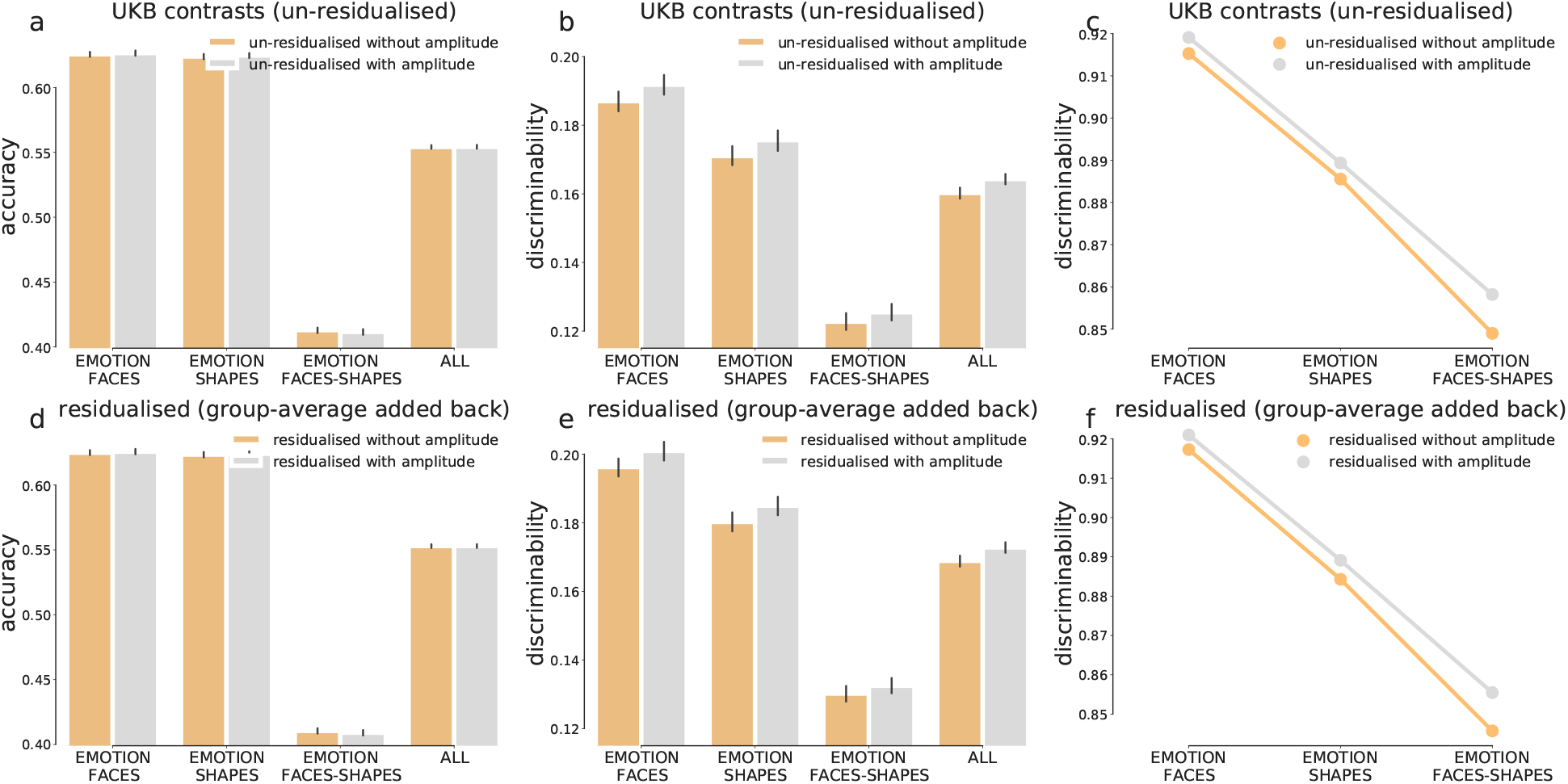
Prediction accuracy and discrimininability of the ensemble model with or without PFM amplitude as additional features, calculated across 17,560 UKB subjects. Although incorporating amplitude did not further increase the overall accuracy for UKB, it did marginally improve prediction discriminability. This coincides with (c) and (d), which shows that the std. maps of predicted activations (across subjects) exhibited higher correspondence with the actual inter-individual variability.

**Figure S14.**
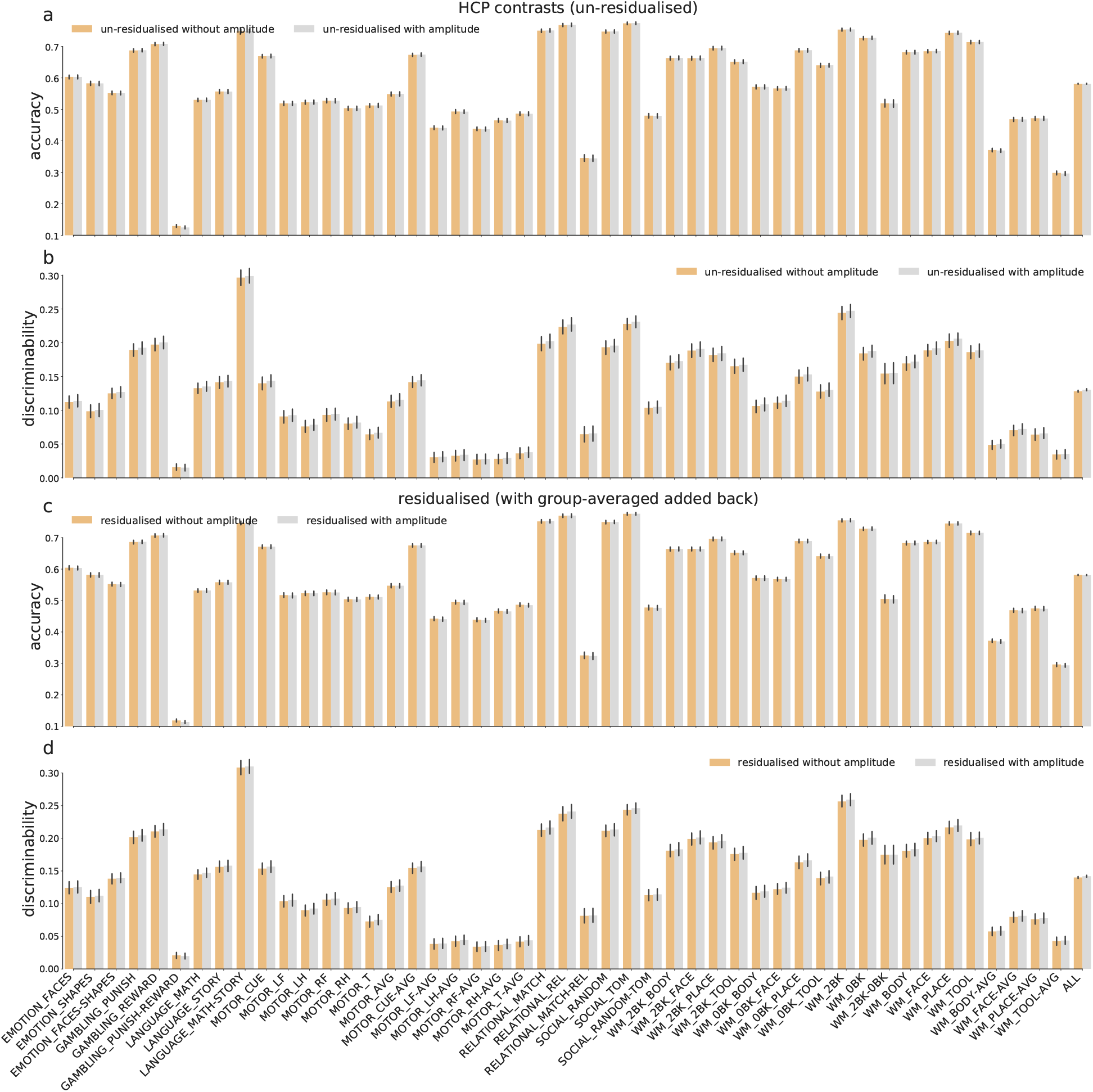
Prediction accuracy and discrimininability of the ensemble model with and without resting-state amplitude as additional features for all 47 HCP contrasts. For HCP, however, including PFM amplitude as additional features at the ensemble stage did not further improve prediction discriminability.

**Figure S15.**
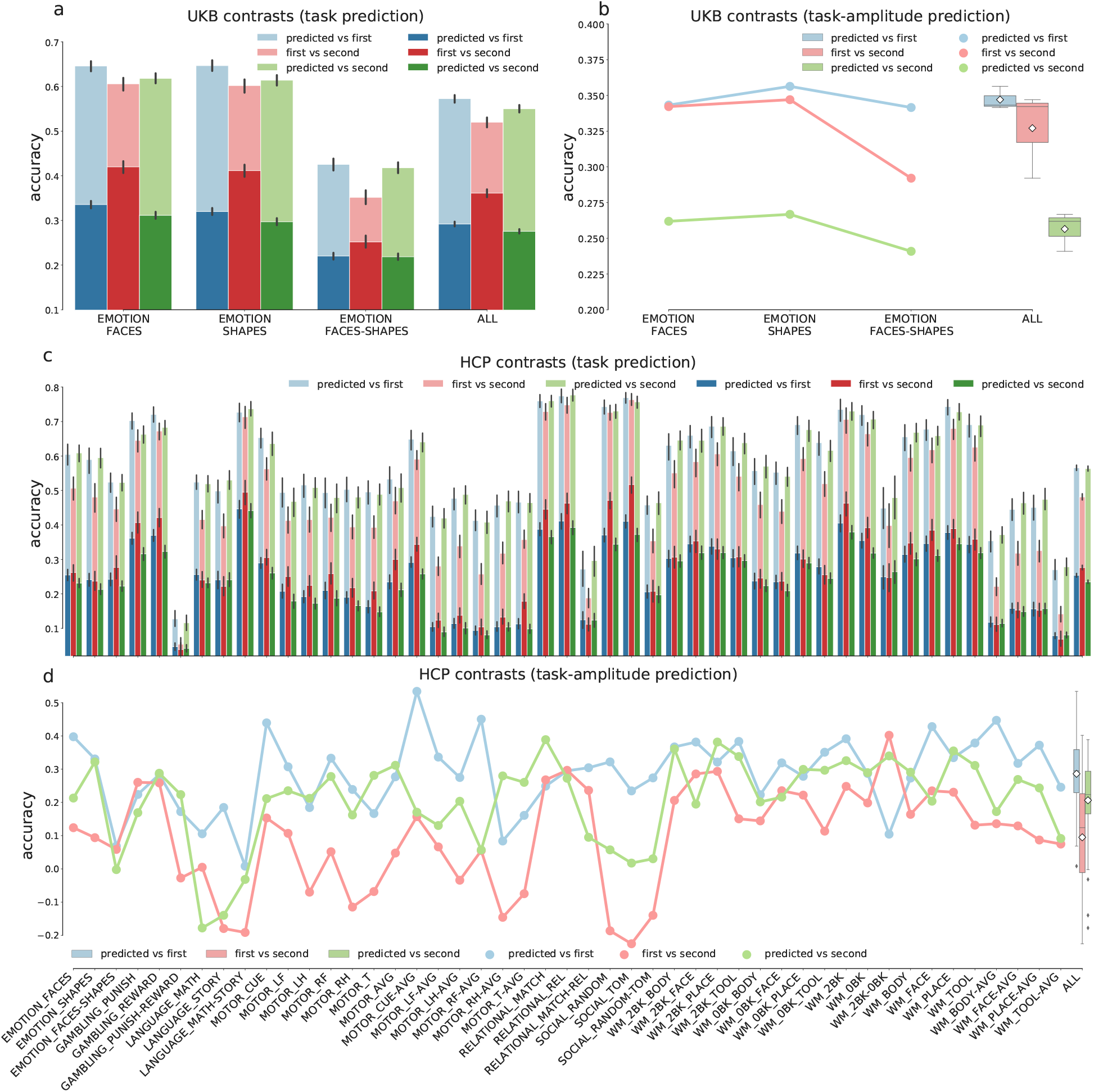
Test-retest reliability of PFM-predicted task maps. The red and blue bars/lines is identical to those shown in Figure 4. The green bars are correlations (accuracy) between the first-time-predicted task maps and the second-visit task contrast maps (note that the second-time data is entirely invisible to training). That the red and green had comparable accuracy suggests the PFM-predicted activations did not overfit to the first-time task-fMRI and generalised well to task-fMRI collected at different visits.

**Figure S16.**
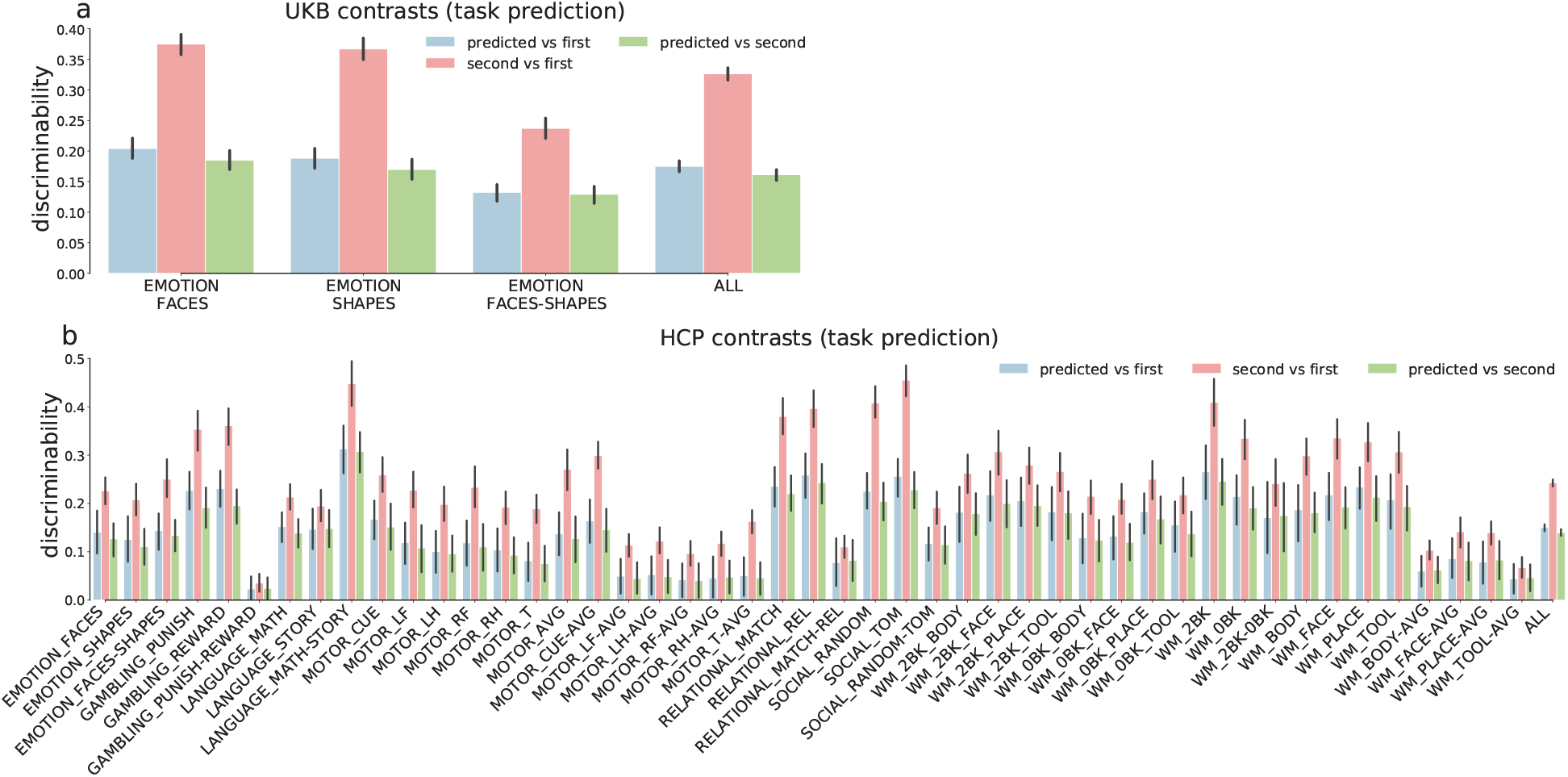
Test-retest reliability of prediction discriminability. Due to the dominance of group-average activations, the predicted task maps (with group-average added back in) yielded lower discriminability than the retest scans.

**Figure S17.**
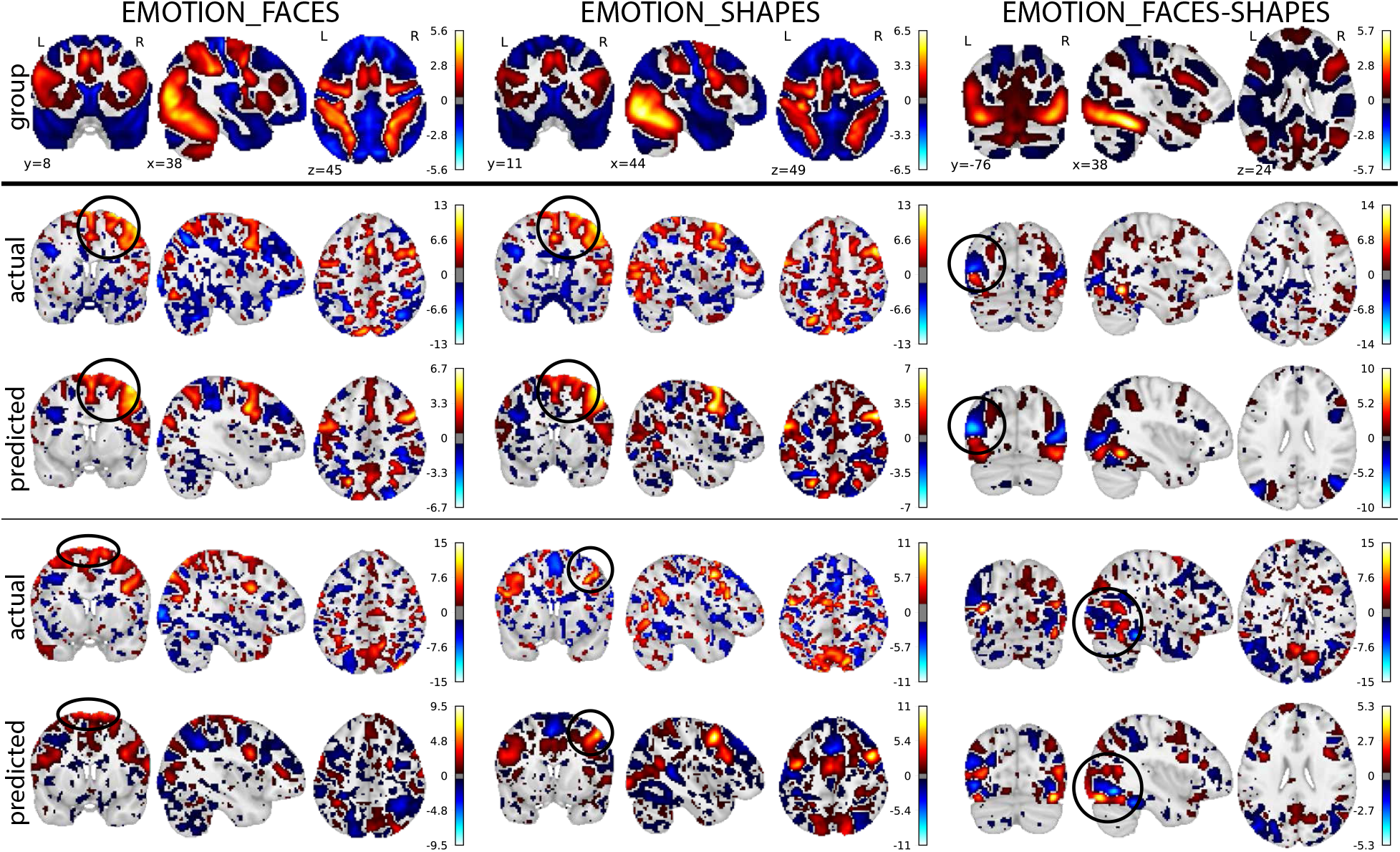
The actual and the predicted task variations (residuals) of the example UKB subjects, shown on the brain.

**Figure S18.**
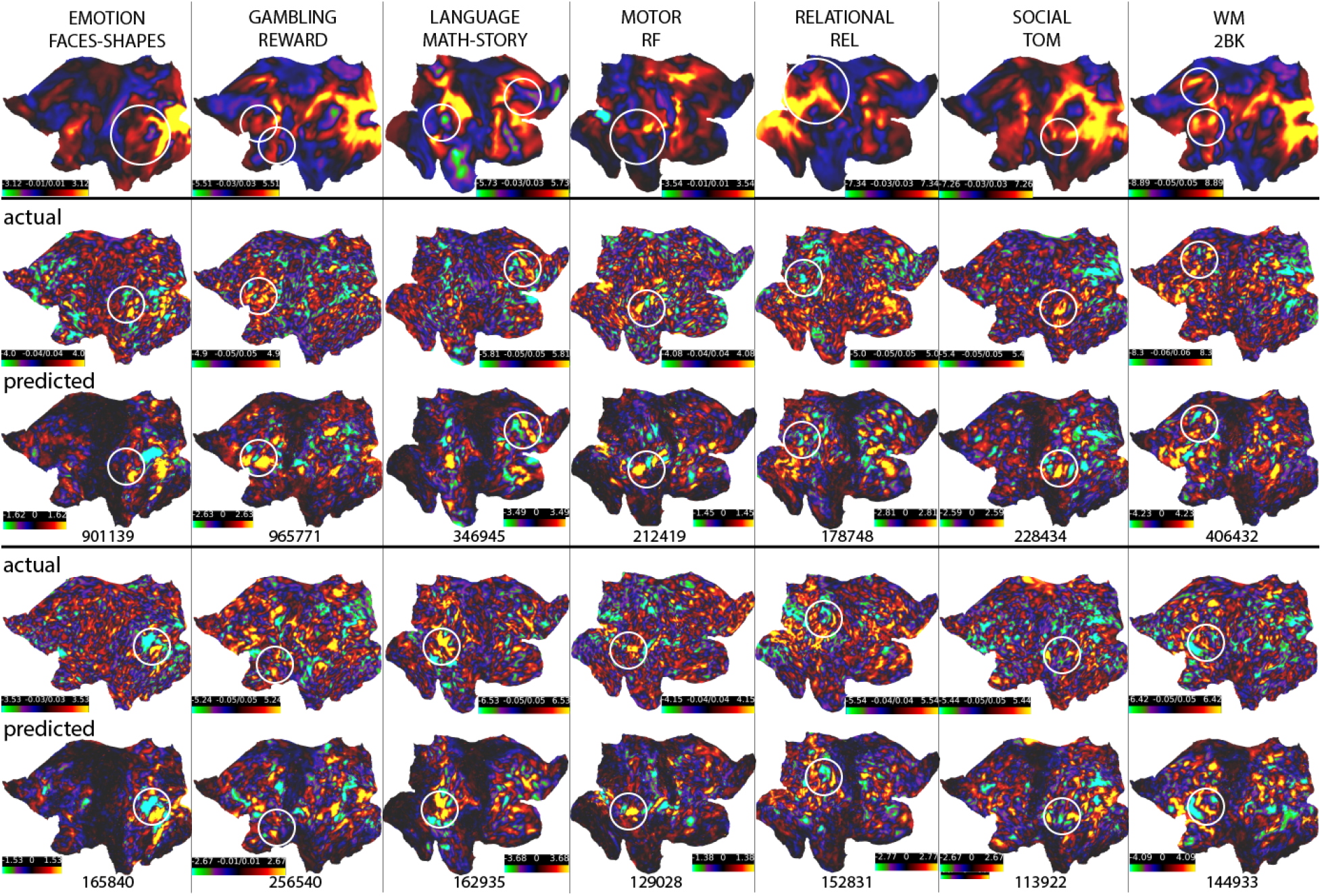
The actual and the predicted task variations (residuals) of the HCP subjects, shown on the surface.

